# Network Medicine Framework Shows Proximity of Polyphenol Targets and Disease Proteins is Predictive of the Therapeutic Effects of Polyphenols

**DOI:** 10.1101/2020.08.27.270173

**Authors:** Italo F. do Valle, Harvey G. Roweth, Michael W. Malloy, Sofia Moco, Denis Barron, Elisabeth Battinelli, Joseph Loscalzo, Albert-László Barabási

## Abstract

Polyphenols, natural products present in plant-based foods, play a protective role against several complex diseases through their antioxidant activity and by diverse molecular mechanisms. Here we developed a network medicine framework to uncover the mechanistic roles of polyphenols on health by considering the molecular interactions between polyphenol protein targets and proteins associated with diseases. We find that the protein targets of polyphenols cluster in specific neighborhoods of the human interactome, whose network proximity to disease proteins is predictive of the molecule’s known therapeutic effects. The methodology recovers known associations, such as the effect of epigallocatechin 3-O-gallate on type 2 diabetes, and predicts that rosmarinic acid (RA) has a direct impact on platelet function, representing a novel mechanism through which it could affect cardiovascular health. We experimentally confirm that RA inhibits platelet aggregation and alpha granule secretion through inhibition of protein tyrosine phosphorylation, offering direct support for the predicted molecular mechanism. Our framework represents a starting point for mechanistic interpretation of the health effects underlying food-related compounds, allowing us to integrate into a predictive framework knowledge on food metabolism, bioavailability, and drug interaction.

## Introduction

Diet plays a defining role in human health. Indeed, while poor diet can significantly increase the risk for coronary heart disease (CHD) and type 2 diabetes mellitus (T2D), a healthy diet can play a protective role, even mitigating genetic risk for CHD^1^. Polyphenols are a class of compounds present in plant-based foods, from fruits to vegetables, nuts, seeds, beans (e.g. coffee, cocoa), herbs, spices, tea, and wine, with well documented protective role as antioxidants, which affect several diseases, from cancer to T2D, cardiovascular, and neurodegenerative diseases^2,3^. Previous efforts profiled over 500 polyphenols in more than 400 foods^4,5^ and have documented the high diversity of polyphenols to which humans are exposed through their diet, ranging from flavonoids to phenolic acids, lignans, and stilbenes.

The underlying molecular mechanisms through which specific polyphenols exert their beneficial effects on human health remain largely unexplored. From a mechanistic perspective, dietary polyphenols are not engaged in endogenous metabolic processes of anabolism and catabolism, but rather affect human health through their anti- or pro-oxidant activity^6^, by binding to proteins and modulating their activity^7,8^, interacting with digestive enzymes^9^, and modulating gut microbiota growth^10,11^. Yet, the variety of experimental settings and the limited scope of studies that explore the molecular effects of polyphenols have, to date, offered a range of often conflicting evidence. For example, two clinical trials, both limited in terms of the number of subjects and the intervention periods, resulted in conflicting conclusions about the beneficial effects of resveratrol on glycemic control in T2D patients^12,13^. We, therefore, need a framework to interpret the evidence present in the literature, and to offer in-depth mechanistic predictions on the molecular pathways responsible for the health implications of polyphenols present in diet. Ultimately, these insights could help us provide evidence on causal diet-health associations, guidelines of food consumption for different individuals, and help to develop novel diagnostic and therapeutic strategies, which may lead to the synthesis of novel drugs.

Here, we address this challenge by developing a network medicine framework to capture the molecular interactions between polyphenols and their cellular binding targets, unveiling their relationship to complex diseases. The developed framework is based on the human interactome, a comprehensive subcellular network consisting of all known physical interactions between human proteins, which has been validated previously as a platform for understanding disease mechanisms^14,15^, rational drug target identification, and drug repurposing^16,17^.

We find that the proteins to which polyphenols bind form identifiable neighborhoods in the human interactome, allowing us to demonstrate that the proximity between polyphenol targets and proteins associated with specific diseases is predictive of the known therapeutic effects of polyphenols. Finally, we unveil the potential therapeutic effects of rosmarinic acid (RA) on vascular diseases (V), predicting that its mechanism of action is related to modulation of platelet function. We confirm this prediction by experiments that indicate that RA modulates platelet function *in vitro* by inhibiting tyrosine protein phosphorylation. Altogether, our results demonstrate that the network-based relationship between disease proteins and polyphenol targets offers a tool to systematically unveil the health effects of polyphenols.

## Results

### Polyphenol Targets Cluster in Specific Functional Neighborhoods of the Interactome

We mapped the targets of 65 polyphenols (see Methods) to the human interactome, consisting of 17,651 proteins and 351,393 interactions (Fig 1a,b). We find that 19 of the 65 polyphenols have only one protein target, while a few polyphenols have an exceptional number of targets (Fig 1c). We computed the Jaccard Index (JI) of the protein targets of each polyphenol pair, finding only a limited similarity of targets among different polyphenols (average JI = 0.0206) (Supplementary Figure 1a). Even though the average JI is small, it is still significantly higher (Z = 147, Supplementary Figure 1b) than the JI expected if the polyphenol targets were randomly assigned from the pool of all network proteins with degrees matching the original set. This finding suggests that while each polyphenol targets a specific set of proteins, their targets are confined to a common pool of proteins, likely determined by commonalities in the polyphenol binding domains of the three-dimensional structure of the protein targets^18^. Gene Ontology (GO) Enrichment Analysis recovers existing mechanisms^8^ and also helps identify new processes related to polyphenol protein targets, such as post-translation protein modifications, regulation, and xenobiotic metabolism (Fig 1d). The enriched GO categories indicate that polyphenols modulate common regulatory processes, but the low similarity in their protein targets, illustrated by the low average JI, indicates that they target different processes within the same process.

**Figure 1.**
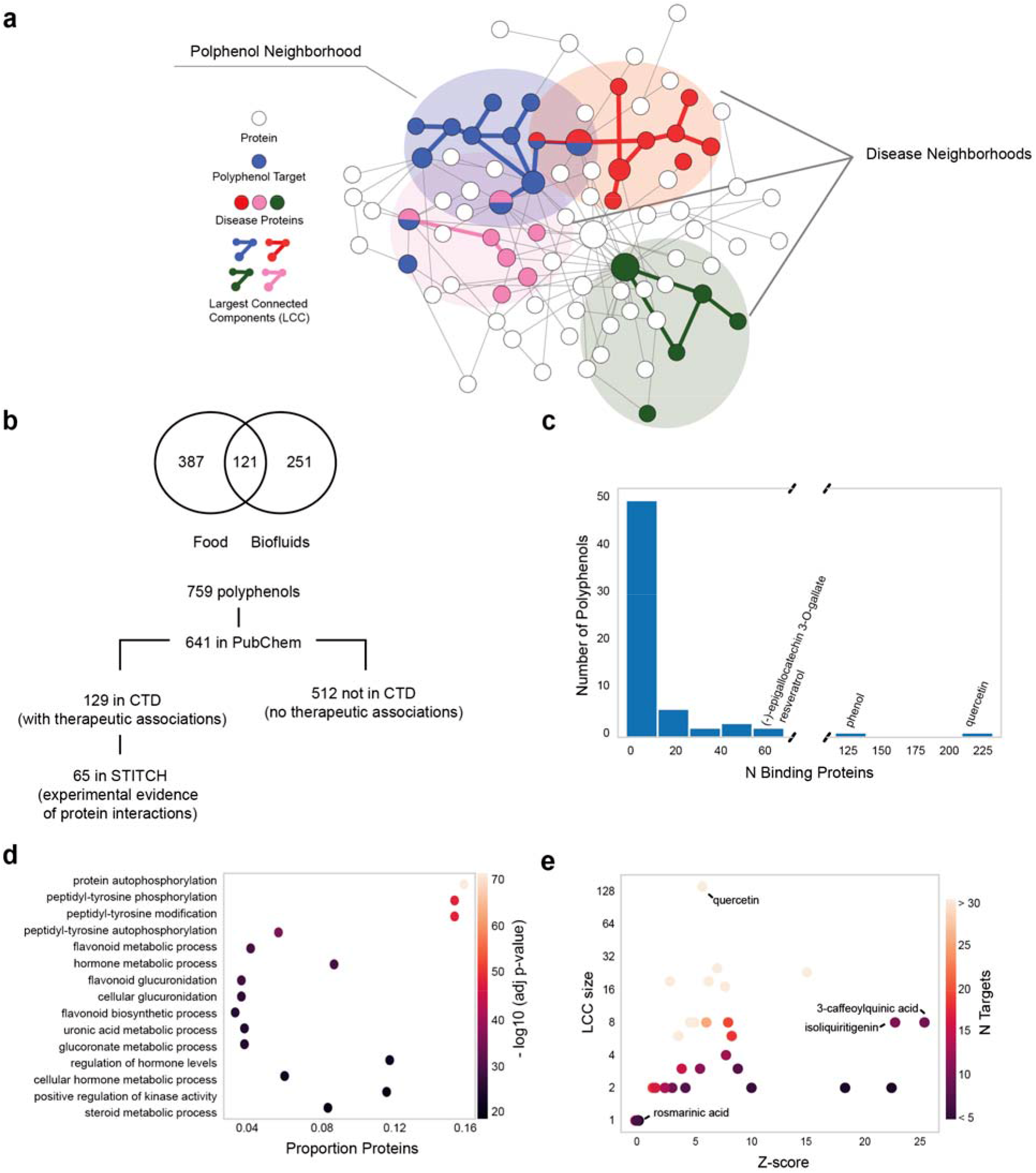
Properties of Polyphenol Protein Targets. (A) Schematic representation of the human interactome, highlighting regions where polyphenol targets and disease proteins are localized. (B) Diagram showing the selection criteria of the polyphenols evaluated in this study. (C) Distribution of the number of polyphenol protein targets mapped to the human interactome. (D) Top (n=15) enriched GO terms (Biological Process) among all polyphenol protein targets. The X-axis shows the proportion of targets mapped to each pathway. (E) Size of the Largest Connected Component (LCC) formed by the targets of each polyphenol in the interactome and the corresponding significance (z-score).

We next asked whether the polyphenol targets cluster in specific regions of the human interactome. We focused on polyphenols with more than two targets (n=46, Fig 2), and measured the size and significance of the largest connected component (LCC) formed by the targets of each polyphenol. We found that 25 of the 46 polyphenols have a larger LCC than expected by chance (Z-score > 1.95) (Fig 1e, Fig 2). In agreement with experimental evidence documenting the effect of polyphenols on multiple pathways^19^, we find that ten polyphenols have their targets organized in multiple connected components of size > 2.

**Figure 2.**
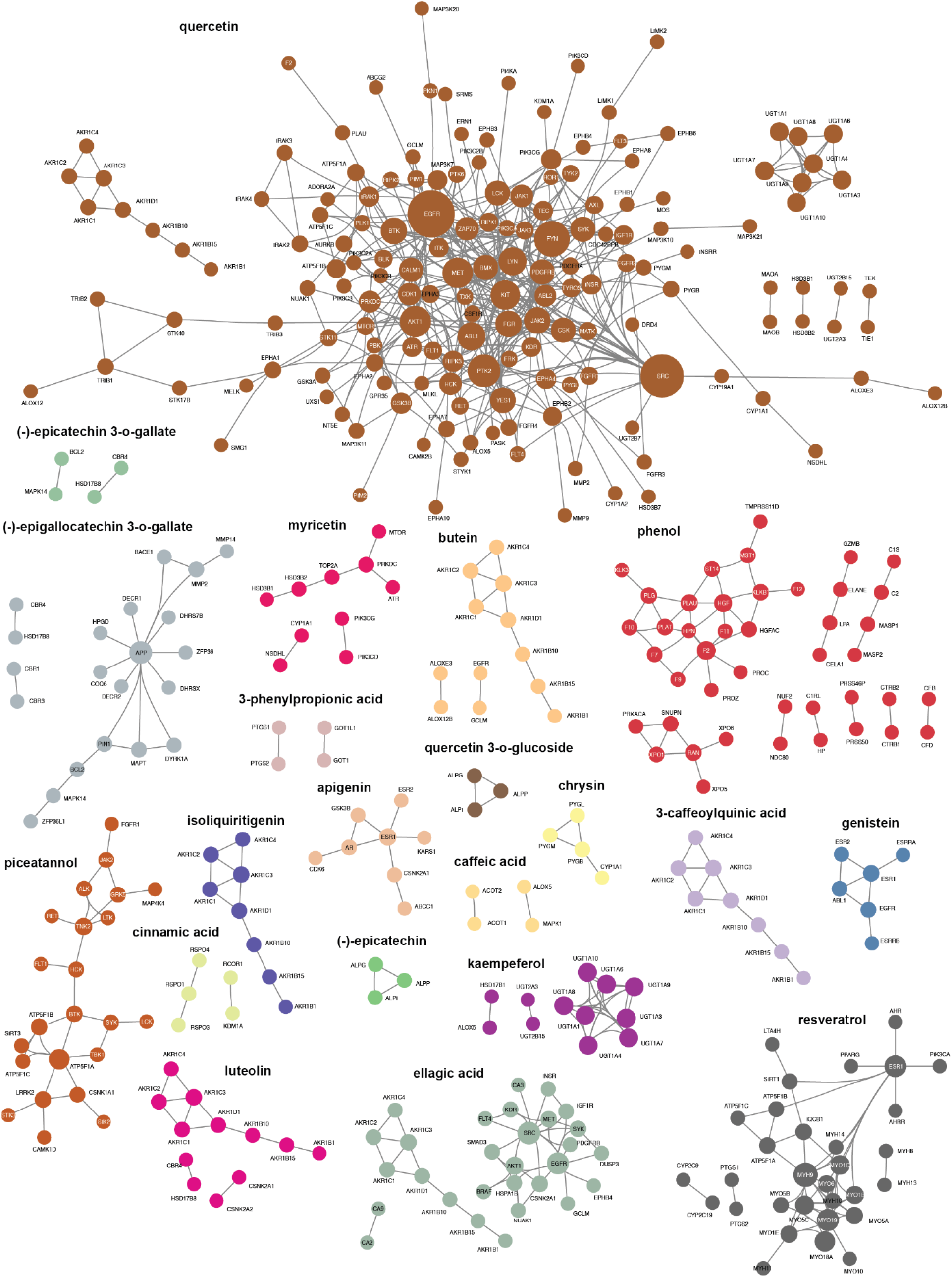
Protein-Protein Interactions of Polyphenol Targets. The 23 polyphenols whose targets form connected components in the interactome and their respective subgraphs. For example, piceatannol targets form a unique connected component of 23 proteins, while quercetin targets form multiple connected components, the largest with 140 proteins. Polyphenol targets that are not connected to any other target are not shown in the figure. Colors distinguish connected component of different polyphenols.

These results indicate that the targets of polyphenols modulate specific well localized neighborhoods of the interactome (Fig 2, Supplementary Figure 1c). This prompted us to explore if the interactome regions targeted by the polyphenols reside within network neighborhoods associated with specific diseases, seeking a network-based framework to unveil the molecular mechanism through which specific polyphenols modulate health.

### Proximity Between Polyphenol Targets and Disease Proteins Reveals their Therapeutic Effects

Polyphenols can be viewed as drugs in that they bind to specific proteins, affecting their ability to perform their normal functions. We, therefore, hypothesized that we can apply the network-based framework used to predict the efficacy of drugs in specific diseases^16,17^ to also predict the therapeutic effects of polyphenols. The closer the targets of a polyphenol are to disease proteins, the more likely that the polyphenol will affect the disease phenotype. We, therefore, calculated the network proximity between polyphenol targets and proteins associated with 299 diseases using the closest measure, *d_c_*, representing the average shortest path length between each polyphenol target and the nearest disease protein (see Methods). Consider for example (−)-epigallocatechin 3-O-gallate (EGCG), a polyphenol abundant in green tea. Epidemiological studies have found a positive relationship between green tea consumption and reduced risk of T2D^20,21^, and physiological and biochemical studies have shown that EGCG presents glucose-lowering effects in both *in vitro* and *in vivo* models^22,23^. We identified 54 experimentally validated EGCG protein targets and mapped them to the interactome, finding that the ECGC targets form an LCC of 17 proteins (Z = 7.61) (Fig 3a). We also computed the network-based distance between EGCG targets and 83 proteins associated with T2D, finding that the two sets are significantly proximal to each other. We ranked all 299 diseases based on the network proximity to the ECGC targets in order to determine whether we can recover the 82 diseases in which ECGC has known therapeutic effects according to the CTD database. By this analysis, we were able to recover 15 previously known therapeutic associations among the top 20 ranked diseases (Table 1), confirming that network-proximity can discriminate between known and unknown disease associations for polyphenols, as previously confirmed among drugs^16,17^.

**Figure 3.**
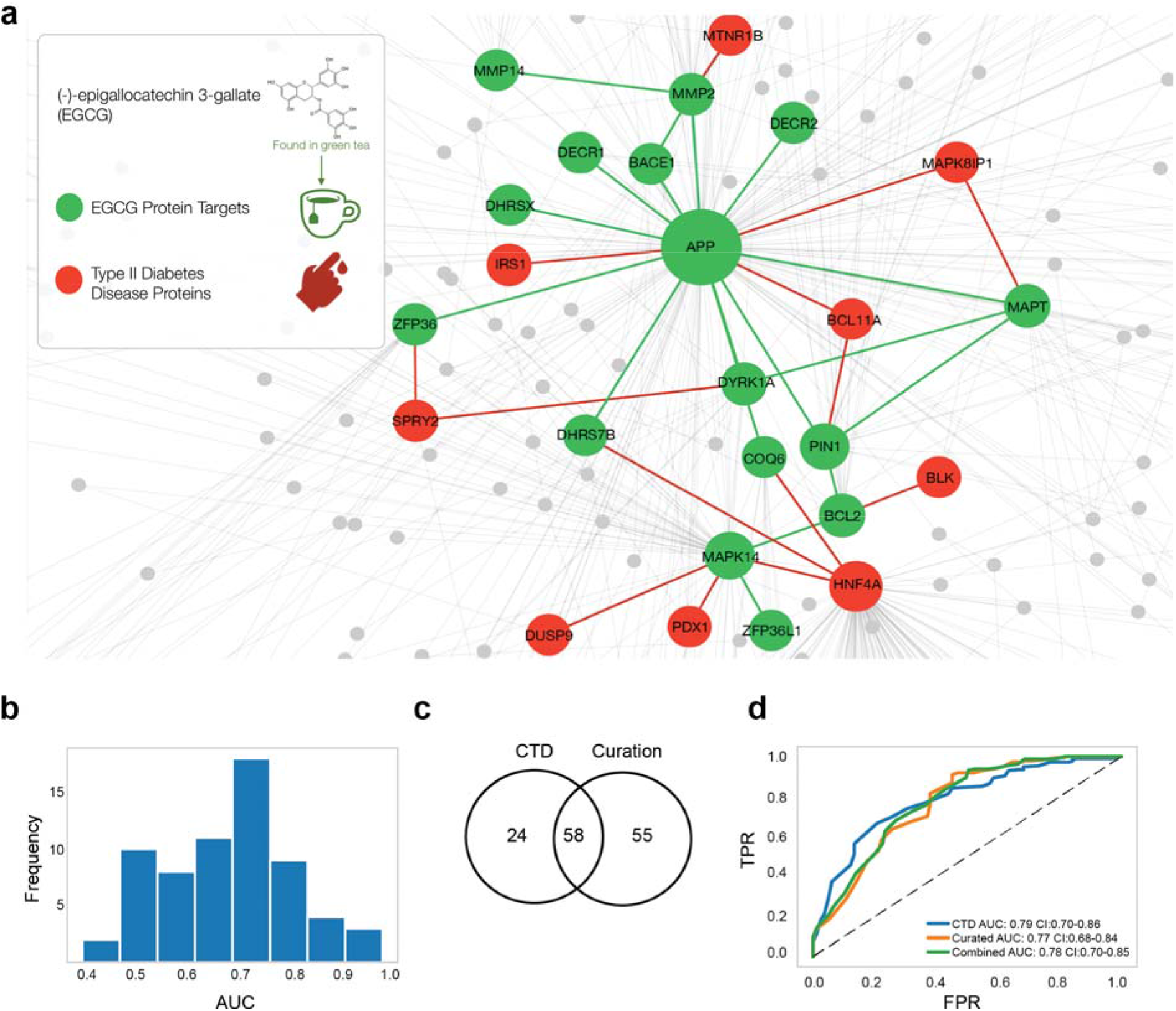
Proximity Between Polyphenol Targets and Disease Proteins is Predictive of Therapeutic Effects of the Polyphenol. (a) Interactome neighborhood showing the EGCG protein targets and their interactions with type 2 diabetes (T2D)-associated proteins. (b) Distribution of AUC values considering the predictions of therapeutic effects for 65 polyphenols. (c) Comparison of the ECGC-disease associations considering the CTD database and the in-house database derived from the manual curation of the literature. (d) Comparison of the prediction performance when considering known EGCG-disease associations from the CTD, in-house manually curated database, or combined datasets.

**Table 1.** Top 20 Predicted Therapeutic Associations Between EGCG and Human Diseases. Diseases were ordered according to the network distance (*d_c_*) of their proteins to EGCG targets and diseases with relative distance *Z_dc_* > −0.5 were removed. References reported in CTD for curated ‘therapeutic associations’ are shown.

| Disease | Distance $d_c$ | Significance $Z_{dc}$ | Known Therapeutic Effect (References) |
| --- | --- | --- | --- |
| nervous system diseases | 1.13 | -1.72 | 64,65 |
| nutritional and metabolic diseases | 1.25 | -1.45 | 23 |
| metabolic diseases | 1.25 | -1.41 | 23 |
| cardiovascular diseases | 1.27 | -2.67 | 66–71 |
| immune system diseases | 1.29 | -1.31 | 72 |
| vascular diseases | 1.33 | -3.47 | 66,67,70 |
| digestive system diseases | 1.33 | -1.57 | 73–77 |
| neurodegenerative diseases | 1.37 | -1.71 | 78 |
| central nervous system diseases | 1.41 | -0.54 | 78 |
| autoimmune diseases | 1.41 | -1.30 | 72 |
| gastrointestinal diseases | 1.43 | -1.02 | 79 |
| brain diseases | 1.43 | -0.89 | NA |
| intestinal diseases | 1.49 | -1.08 | 79 |
| inflammatory bowel diseases | 1.54 | -2.10 | NA |
| bone diseases | 1.54 | -1.18 | NA |
| gastroenteritis | 1.54 | -1.92 | NA |
| demyelinating diseases | 1.54 | -1.78 | NA |
| glucose metabolism disorders | 1.54 | -1.58 | 23 |
| heart diseases | 1.56 | -1.20 | 68,69,71 |
| diabetes mellitus | 1.56 | -1.66 | 23 |

We expanded these methods to all polyphenol-disease pairs, to predict diseases for which specific polyphenols might have therapeutic effects. For this analysis, we grouped all 19,435 polyphenol-disease associations between 65 polyphenols and 299 diseases into known (1,525) and unknown (17,910) associations. The known polyphenol-disease set was retrieved from CTD, which is limited to manually curated associations for which there is literature-based evidence. For each polyphenol, we tested how well network proximity discriminates between the known and unknown sets by evaluating the area under the Receiving Operating Characteristic (ROC) curve (AUC). For EGCG, network proximity offers good discriminative power (AUC = 0.78, CI: 0.70 - 0.86) between diseases with known and unknown therapeutic associations (Table 1). We find that network proximity (*d_c_*) offers predictive power with an AUC > 0.7 for 31 polyphenols (Fig 3b). The methodology recovers many associations well documented in the literature, like the beneficial effects of umbelliferone on colonrectal neoplasms^24,25^.

In Table 2 we summarize the top 10 polyphenols for which the network medicine framework offers the best predictive power of therapeutic effects, limiting the entries to predictive performance of AUC > 0.6 and performance over top predictions with precision > 0.6. Given the lack of data on true negative examples, we considered unknown associations as negative cases, observing the same trend when we used an alternative performance metric that does not require true negative labels (i.e. AUC of the Precision-Recall curve) (Supplementary Figure 2).

**Table 2.** Top Ranked Polyphenols. Polyphenols for which network proximity to diseases best predicts their therapeutic effects. Table showing polyphenols with AUC > 0.6 and Precision > 0.6. (*) Confidence intervals calculated with 2,000 bootstraps with replacement and sample size of 50% of the diseases (150/299). (**) Precision was calculated based on the top 10 polyphenols after their ranking based on the distance (d_c_) of their targets to the disease proteins and considering only predictions with Z-score < −0.5.(***) Concentrations of polyphenols in blood were retrieved from the Human Metabolome Database (HMDB)

| Polyphenol | AUC | AUC CI* | Precision** | Concentration in Blood*** | N Mapped Targets | LCC Size |
| --- | --- | --- | --- | --- | --- | --- |
| Coumarin | 0.93 | [0.86 - 0.98] | 0.6 |  | 7 | 1 |
| Piceatannol | 0.86 | [0.77 - 0.94] | 0.6 |  | 39 | 23 |
| Genistein | 0.82 | [0.75 - 0.89] | 0.7 | [0.006 - 0.525 uM] | 18 | 6 |
| Ellagic acid | 0.79 | [0.63 - 0.92] | 0.6 |  | 42 | 19 |
| (-)-epigallocatechin 3-O-gallate | 0.78 | [0.70 - 0.86] | 0.8 |  | 51 | 17 |
| Isoliquiritigenin | 0.75 | [0.77 - 0.94] | 0.6 |  | 10 | 8 |
| Resveratrol | 0.75 | [0.66 - 0.82] | 1 |  | 63 | 25 |
| Pterostilbene | 0.73 | [0.61 - 0.84] | 0.6 |  | 5 | 2 |
| Quercetin | 0.73 | [0.64 - 0.81] | 1 | [0.022 - 0.080 uM] | 216 | 140 |
| (-)-epicatechin | 0.65 | [0.49 - 0.80] | 0.8 | 0.625 uM | 11 | 3 |

Finally, we performed multiple robustness checks to exclude the role of potential biases in the input data. To test if the predictions are biased by the set of known associations retrieved from CTD, we randomly selected 100 papers from PubMed containing MeSH terms that tag EGCG to diseases. We manually curated the evidence for EGCG’s therapeutic effects for the diseases discussed in the published papers, excluding reviews and non-English language publications. The dataset was processed to include implicit associations (see Methods), resulting in a total of 113 diseases associated with EGCG, of which 58 overlap with the associations reported by CTD (Fig 3c). We observed that the predictive power of network proximity was unaffected by whether we considered the annotations from CTD, the manually curated list, or the union of both (Fig 3d). To test the role of potential biases in the interactome, we repeated our analysis using only high-quality polyphenol-protein interactions retrieved from ligand-protein 3D resolved structures (Supplementary Figure 1d) and a subset of the interactome derived from an unbiased high-throughput screening (Supplementary Figure 1f). We find that the predictive power was largely unchanged, indicating that the literature bias in the interactome does not affect our findings. Finally, we re-tested the predictive performance by considering not only the therapeutic polyphenol-disease associations, but also the marker/mechanism ones - another type of curated association available in CTD - finding that the predictive power remains largely unchanged (Supplementary Notes, Supplementary Figure 3).

### Network Proximity Predicts Gene Expression Perturbation Induced by Polyphenols

To validate that network proximity reflects biological activity of polyphenols observed in experimental data, we retrieved expression perturbation signatures from the Connectivity Map database^26^ for the treatment of the breast cancer MCF7 cell line with 21 polyphenols (Supplementary Table 1, Supplementary Figure 4). We investigated the relationship between the extent to which polyphenols perturb the expression of disease genes, the network proximity between the polyphenol targets and disease proteins, and their known therapeutic effects (Fig 4a). For example, we observe different perturbation profiles for gene pools associated with different diseases: for treatment with genistein (1 μM, 6 hours) we observe 10 skin disease genes with perturbation score > 2, while we observe only one highly perturbed cerebrovascular disorder gene (Fig 4b). Indeed, network proximity indicates that skin disease is closer to the genistein targets than cerebrovascular disorder, suggesting a relationship between network proximity, gene expression perturbation, and the therapeutic effects of the polyphenol (Fig 4a). To test this hypothesis, we computed an enrichment score that measures the overrepresentation of disease genes among the most perturbed genes (see Methods), finding 13 diseases that have their genes significantly enriched among the most deregulated genes by genistein, of which 4 have known therapeutic associations. We find that these four diseases are significantly closer to the genistein targets than the nine diseases with unknown therapeutic associations (Fig 4c). We observed a similar trend for treatments with other polyphenols, whether we use the same (1μM, Fig 4c) or different (100nM to 10μM, Supplementary Figure 5) concentrations. This result suggests that changes in gene expression caused by a polyphenol are indicative of its therapeutic effects, but only if the observed expression change is limited to proteins proximal to the polyphenol targets (Fig 4a).

**Figure 4.**
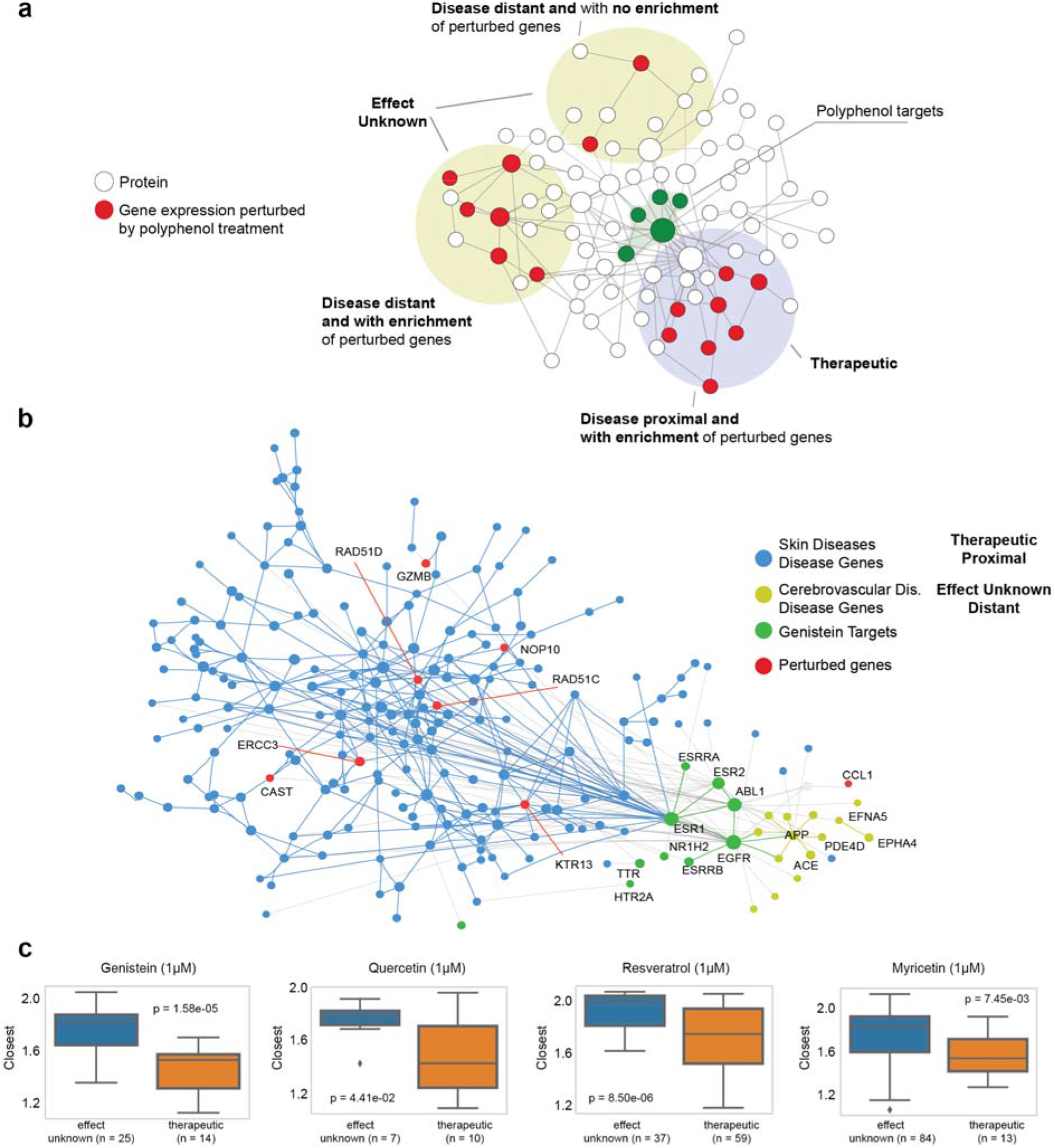
Relationships among Gene Expression Perturbation, Network Proximity, and Therapeutic Effects of Polyphenols on Diseases. (a) Schematic representation of the relationship between the extent to which a polyphenol perturbs disease genes expression, its proximity to the disease genes, and its therapeutic effects. (b) Interactome neighborhood showing the modules of skin diseases (SD), genistein, and cerebrovascular disorders (CD). The SD module has 10 proteins with high perturbation scores (>2) in the treatment of the MCF7 cell line with 1 μM of genistein. Genes associated to SD are significantly enriched among the most differentially expressed genes, and the maximum perturbation score among disease genes is higher in SD than CD. (c) Among the diseases in which genes are enriched with highly perturbed genes, those with therapeutic associations show smaller network distances to the polyphenol targets than those without. The same trend is observed in treatments of the polyphenols quercetin, resveratrol, and myricetin.

Consequently, network proximity should also be predictive of the overall gene expression perturbation caused by a polyphenol on the genes of a given disease. To test this hypothesis, in each experimental combination defined by the polyphenol type and its concentration, we evaluated the maximum perturbation among genes for each disease. We then compared the magnitude of the observed perturbation between diseases that were proximal (*d_c_* < 25^th^ percentile, 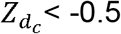) or distal (*d_c_* > 75^th^ percentile, 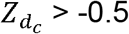) to the polyphenol targets. Figures 5a-b and Supplementary Figure 6 show the results for the genistein treatment (1μM, 6 hours), indicating that diseases proximal to the polyphenol targets show higher maximum perturbation values than distal diseases. The same trend is observed for other polyphenols when we use different *d_c_* and 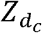 thresholds for defining proximal and distant diseases (Figs 5b, Supplementary Figures 6-9), confirming that the impact of a polyphenol on cellular signaling pathways is localized in the network space, being greater in the vicinity of the polyphenol targets compared to neighborhoods remote from these targets. We also considered gene expression perturbations in the network vicinity of the polyphenol targets, regardless of whether the proteins were disease proteins or not, observing higher perturbation scores for proximal proteins in 12 out 21 polyphenols tested at 10μM (Supplementary Figure 10). Finally, we find that the enrichment score of perturbed genes among disease genes is not as predictive of the polyphenol therapeutic effects as network proximity (Supplementary Figure 11).

**Figure 5.**
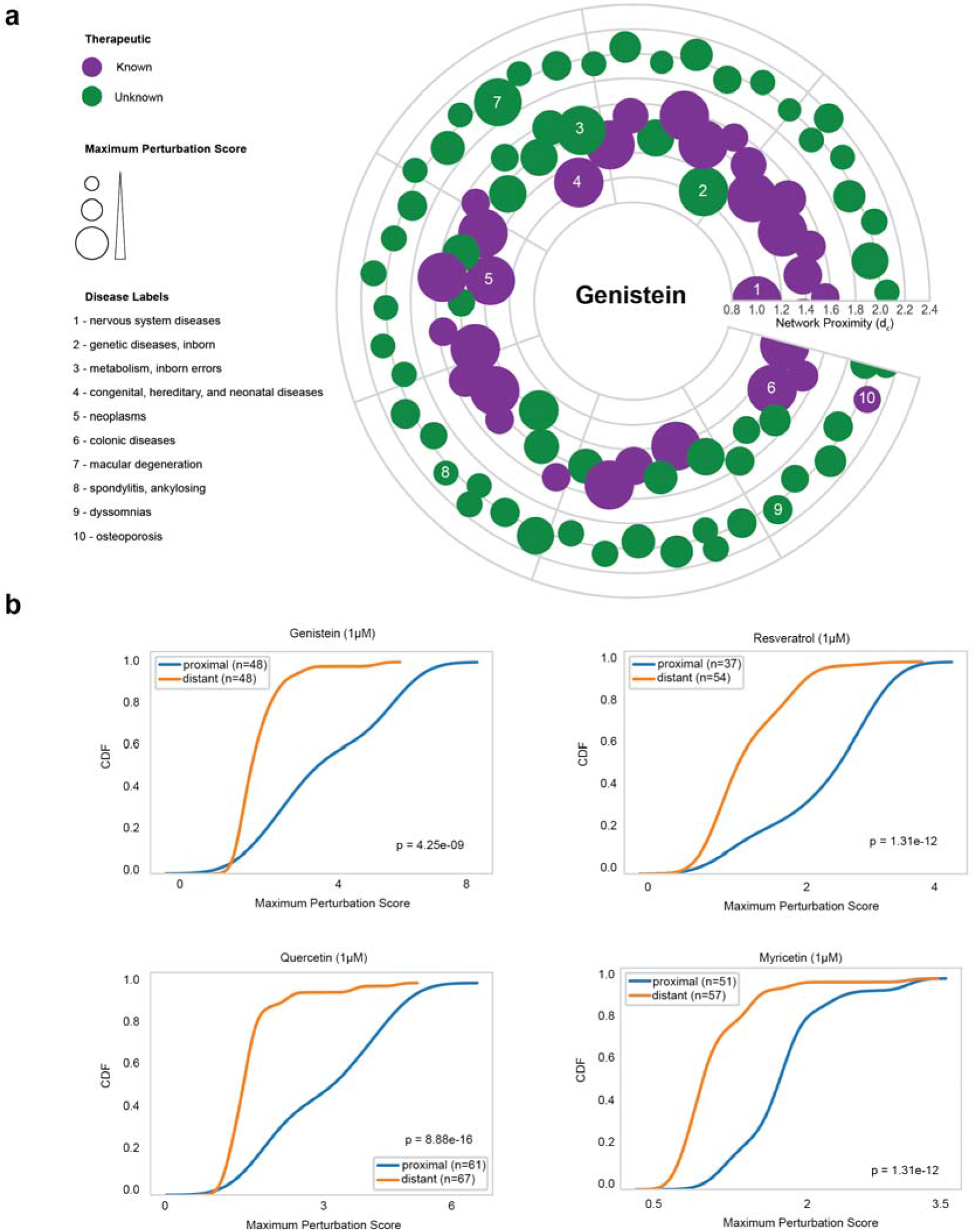
Diseases Proximal to Polyphenol Targets Have Higher Gene Expression Perturbation Profiles. (a) Proximal and distal diseases in relation to genistein targets. Each node represents a disease and the node size is proportional to the perturbation score after treatment with genistein (1 μM, 6 hours). Distance from the origin represents the network proximity (*d_c_*) to genistein targets. Purple nodes represent diseases in which the therapeutic association was previously known. (c) Cumulative distribution of the maximum perturbation scores of genes from diseases that are distal or proximal to polyphenol targets considering different polyphenols (1 μM, 6 hours): genistein, quercetin, resveratrol, and myricetin. Statistical significance was evaluated with the Kolmogorov-Smirnov test.

Altogether these results indicate that network proximity offers a mechanistic interpretation for the gene expression perturbations induced by polyphenols on disease genes. They also show that network proximity can indicate when gene expression perturbations result in therapeutic effects, suggesting that future studies could integrate gene expression (whenever available) with network proximity as they aim to more accurately prioritize polyphenol-disease associations.

### Experimental Evidence Confirms that Rosmarinic Acid Modulates Platelet Function

To demonstrate how the network-based framework can facilitate the mechanistic interpretation of the therapeutic effects of selected polyphenols, we next focus on vascular diseases (V). Of 65 polyphenols evaluated in this study, we found 27 to have associations to V, as their targets were within the V network neighborhood (Supplementary Table 3). We, therefore, inspected the targets of 15 of the 27 polyphenols with 10 or less targets. The network analysis identified direct links between biological processes related to vascular health and the targets of three polyphenols: gallic acid, rosmarinic acid, and 1,4-naphthoquinone (Supplementary Figure 12, Supplementary Notes). The network neighborhood containing the targets of these polyphenols suggests that gallic acid activity involves thrombus dissolution processes, rosmarinic acid acts on platelet activation and antioxidant pathways through FYN and its neighbors, and 1,4-naphthoquinone acts on signaling pathways of vascular cells through MAP2K1 activity (Supplementary Figure 12, Supplementary Notes).

To validate the developed framework, we set out to obtain direct experimental evidence of the predicted mechanistic role of rosmarinic acid (RA) in V. The RA targets are in close proximity to proteins related to platelet function, forming the RA-V-platelet module: a connected component formed by the RA target FYN and the V proteins associated with platelet function PDE4D, CD36, and APP (Fig 6a). We, therefore, asked whether RA influenced platelet activation *in vitro*. As platelets can be stimulated through different activation pathways, RA effects can, in principle, occur in any of them. To test these different possibilities, we pretreated platelets with RA and then activated with: 1) glycoprotein VI by collagen or collagen-related peptide (CRP/CRPXL); 2) protease-activated receptors-1,4 by thrombin receptor activator peptide-6 (TRAP-6); 3) prostanoid thromboxane receptor by the thromboxane A_2_ analogue (U46619); and 4) P2Y1/12 receptor by adenosine diphosphate (ADP)^27^. When we compared the network distance between each stimulant receptor and the RA-V-platelet module (Fig 6a), we observed that the receptors for CRP/CRPXL, TRAP-6, and U46619 are closer than random expectation, while the receptor for ADP is more distant (Fig 6b). We expected that platelets would be most affected by RA when treated with stimulants whose receptors are most proximal to the RA-V-platelet module, i.e., CRP/CRPXL, TRAP-6, and U46619, and as a control, we expect no effect for the distant ADP receptor. The experiments confirm this prediction: RA inhibits collagen-mediated platelet aggregation (Fig 6c) and impairs dense granule secretion induced by CRPXL, TRAP-6, and U46619 (Supplementary Figure 13). RA-treated platelets also displayed dampened alpha-granule secretion (Fig 6d) and integrin αIIbβ3 activation (Supplementary Figure 13) in response to U46619. As expected, RA did not affect platelet function when we used an agonist whose receptor is distant from the RA-V-module, i.e., ADP. These findings suggest that RA impairs basic hallmarks of platelet activation via strong network effects, supporting our hypothesis that the proximity between RA targets and the neighborhood associated with platelet function (Fig 6a) could in part explain RA’s impact on V.

**Figure 6.**
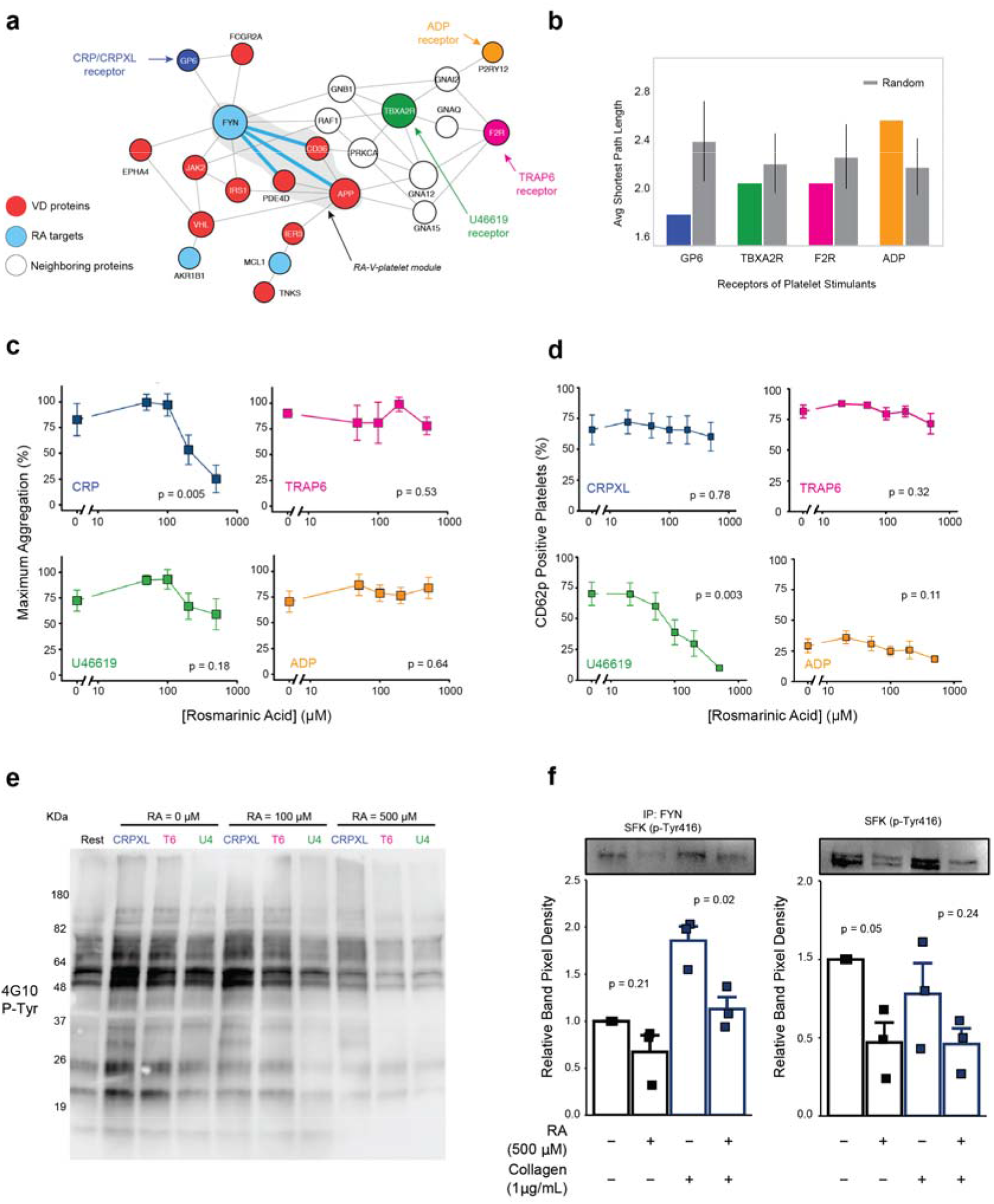
Rosmarinic Acid Modulates Platelet Function. (a) Interactome neighborhood showing rosmarinic acid (RA) targets and the RA-V-platelet module - the connected component formed by the RA target FYN and the V proteins associated with platelet function PDE4D, CD36, and APP – and the receptor for platelet agonists used in our experiments (collagen/CRPXL, TRAP6, U46619, and ADP). (b) Average shortest path length from each platelet agonist receptor and the RA-V-platelet module formed by the proteins FYN, PDE4D, CD36, APP. Bars represent standard deviation of that same measure over 1000 iterations of random selection of nodes in a degree preserving fashion. c-e) Platelet-rich plasma (PRP) or washed platelets were pre-treated with RA for 1 hour before stimulation with either collagen (1 μg/mL), collagen-related peptide (CRP-XL, 1μg/mL), thrombin receptor activator peptide-6 (TRAP-6, 20 μM), U46619 (1 μM), or ADP (10 μM). Platelets were assessed for either (c) aggregation, (d) alpha granule secretion. Platelet lysates were also probed for either (e) non-specific tyrosine phosphorylation (p-Tyr) of the whole cell lysate, or (d) site-specific phosphorylation of src family kinases (SFKs) and FYN at residue 416. n = 3-6 separate blood donations, mean +/− SEM. p-values in (c) and (d) were determined by Kruskal-Wallis test and by unpaired t.tests in (f).

We next searched to clarify the molecular mechanisms involved in the impact of RA on platelets. Given that platelet activation is coordinated by several kinases, we hypothesized that RA inhibits platelet function by blocking agonist-induced protein tyrosine phosphorylation. We observed that RA-treated platelets demonstrated a dose-dependent reduction in total tyrosine phosphorylation in response to CRPXL, TRAP-6 and U46619 (Fig 6e). Given that RA caused a substantial decrease in phosphorylation of proteins with atomic mass between 50-60 KDa (Fig 6e), we hypothesized that RA may reduce phosphorylation of FYN (59 KDa), or other similarly sized members of the same protein family (*i.e.* src family kinases, SFKs). To test this, we measured the level of phosphorylation within the activation domain (amino acid 416) of SFKs, finding that RA reduced collagen induced phosphorylation of FYN as well as basal tyrosine phosphorylation of SFKs (Fig 6f). This indicates that RA perturbs the phospho-signaling networks that regulate platelet response to extracellular stimuli.

Altogether, these findings support our prediction that RA modulates platelet activation and function. It also supports the observation that its mechanism of action involves reduction of phosphorylation at the activation domain of the protein-tyrosine kinase FYN (Fig 6a) and the inhibition of general tyrosine phosphorylation. Finally, while polyphenols are usually associated to their antioxidant function, here we illustrate another mechanistic pathway through which they could benefit health.

## Discussion

Here, we proposed a network-based framework to predict the therapeutic effects of dietary polyphenols in human diseases. We find that polyphenol protein targets cluster in specific functional neighborhoods of the interactome, and we show that the network proximity between polyphenol targets and disease proteins is predictive of the therapeutic effects of polyphenols. We demonstrate that diseases whose proteins are proximal to polyphenol targets tend to have significant changes in gene expression in cell lines treated with the respective polyphenol, while such changes are absent for diseases whose proteins are distal to polyphenol targets. Finally, we find that the network neighborhood around the RA targets and vascular disease proteins are related to platelet function. We validate this mechanistic prediction by showing that RA modulates platelet function through inhibition of protein tyrosine phosphorylation. These observations suggest a role of RA on prevention of vascular diseases by inhibiting platelet activation and aggregation.

The observed results also suggest multiple avenues through which our ability to understand the role of polyphenols could be improved. First, some of the known health benefits of polyphenols might be caused not only by the native molecules, but also by their metabolic byproducts ^28,29^. We, however, lack data about colonic degradation, liver metabolism, bioavailability, and interaction with proteins of specific polyphenols or their metabolic byproducts. Future experimental data on protein interactions with polyphenol byproducts and conjugates can be incorporated in the proposed framework, further improving the accuracy of our predictions. The lack of this data does not invalidate the findings presented here, since previous studies report the presence of unmetabolized polyphenols in blood^30–32^; and it has been hypothesized that, in some instances, deconjugation of liver metabolites occurs in specific tissues or cells^33–35^. Therefore, the lack of data for specific polyphenols and the fact that other mechanisms exist through which they can affect health (e.g. antioxidant activity, microbiota regulation) explain why this methodology might still miss a few known relationships between polyphenols and diseases. Second, considering that several experimental studies of polyphenol bioefficacy have been observed in *in vitro* and *in vivo* models, the proposed framework might help us interpret literature evidence, possibly even allowing us to exclude chemical candidates when considering the health benefits provided by a given food in epidemiological association studies.

Our assumption that network proximity recovers therapeutic associations is based on its predictive performance on a ground truth dataset for observed therapeutic effects and also relies on previous observations about the effect of drugs on diseases^16,17,36^. While the proposed methodology offers a powerful prioritization tool to guide future research, the real effect of polyphenols on diseases might still be negative, given other unmet factors such as dosage, comorbidities, and drug interactions, which can only be ruled out by pre-clinical and clinical studies. Gene expression perturbation profiles, such as the ones provided by the Connectivity map, can also be integrated with network proximity to further highlight potential beneficial or harmful effects of chemical compounds^37,38^.

The low bioavailability of some polyphenols in food might still present challenges when considering the therapeutic utility of these molecules. However, 48 of the 65 polyphenols we explored here are predicted to have high gastrointestinal absorption (Supplementary Table 2) and different methodologies are available to increase bioavailability of natural compounds^39,40^. Additionally, in the same way that the polyphenol phlorizin led to the discovery of new strategies for disease treatment resulting in the development of new compounds with higher efficacy^41^, we believe that the present methodology can help us identify polyphenol-based candidates for drug development.

The methodology introduced here offers a foundation for the mechanistic interpretation of alternative pathways through which polyphenols can affect health, e.g., the combined effect of different polyphenols^36,42^ and their interaction with drugs^43^. To address such synergistic effects, we need ground-truth data on these aspects. The developed methodology can be applied to other food-related chemicals, providing a framework by which to understand their health effects. Future research may help us also account for the way that food-related chemicals affect endogenous metabolic reactions, impacting not only signaling pathways, but also catabolic and anabolic processes. Finally, the methodology provides a framework to interpret and find causal support for associations identified in observational studies. Taken together, the proposed network-based framework has the potential to reveal systematically the mechanism of action underlying the health benefits of polyphenols, offering a logical, rational strategy for mechanism-based drug development of food-based compounds.

## Methods

### Building the Interactome

The human interactome was assembled from 16 databases containing different types of protein-protein interactions (PPIs): 1) binary PPIs tested by high-throughput yeast-two-hybrid (Y2H) experiments^44^; 2) kinase-substrate interactions from literature-derived low-throughput and high-throughput experiments from KinomeNetworkX^45^, Human Protein Resource Database (HPRD)^46^, and PhosphositePlus^47^; 3) carefully literature-curated PPIs identified by affinity purification followed by mass spectrometry (AP-MS), and from literature-derived low-throughput experiments from InWeb^48^, BioGRID^49^, PINA^50^, HPRD^51^, MINT^52^, IntAct^52^, and InnateDB^53^; 4) high-quality PPIs from three-dimensional (3D) protein structures reported in Instruct^54^, Interactome3D^55^, and INSIDER^56^; 5) signaling networks from literature-derived low-throughput experiments as annotated in SignaLink2.0^57^; and 6) protein complex from BioPlex2.0^58^. The genes were mapped to their Entrez ID based on the National Center for Biotechnology Information (NCBI) database as well as their official gene symbols. The resulting interactome includes 351,444 protein-protein interactions (PPIs) connecting 17,706 unique proteins (Supplementary Data 1). The largest connected component has 351,393 PPIs and 17,651 proteins.

### Polyphenols, Polyphenol Targets, and Disease Proteins

We retrieved 759 polyphenols from the PhenolExplorer database^4^. The database lists polyphenols with food composition data or profiled in biofluids after interventions with polyphenol-rich diets. For our analysis, we only considered polyphenols that: 1) could be mapped in PubChem IDs, 2) were listed in the Comparative Toxicogenomics (CTD) database^59^ as having therapeutic effects on human diseases, and 3) had protein-binding information present in the STITCH database^60^ with experimental evidence (Fig 1a). After these steps, we considered a final list of 65 polyphenols, for which 598 protein targets were retrieved from STITCH (Supplementary Table 1). We considered 3,173 disease proteins corresponding to 299 diseases retrieved from Menche *et al* (2015)^15^. Gene ontology enrichment analysis of protein targets was performed using the Bioconductor package clusterProfiler with a significance threshold of p < 0.05 and Benjamini-Hochberg multiple testing correction with q < 0.05.

### Polyphenol Disease Associations

We retrieved the polyphenol-disease associations from the Comparative Toxicogenomics Database (CTD). We considered only manually curated associations labeled as therapeutic. By considering the hierarchical structure of diseases along the MeSH tree, we expanded explicit polyphenol-disease associations to include also implicit associations. This procedure was performed by propagating associations in the lower branches of the MeSH tree to consider diseases in the higher levels of the same tree branch. For example, a polyphenol associated with heart diseases would also be associated with the more general category of cardiovascular diseases. By performing this expansion, we obtained a final list of 1,525 known associations between the 65 polyphenols and the 299 diseases considered in this study.

### Network Proximity Between Polyphenol Targets and Disease Proteins

The proximity between a disease and a polyphenol was evaluated using a distance metric that takes into account the shortest path lengths between polyphenol targets and disease proteins^16^. Given *S*, the set of disease proteins, *T*, the set of polyphenol targets, and *d*(*s, t*), the shortest path length between nodes *s* and *t* in the network, we define:

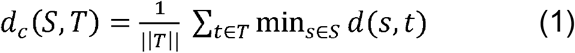

We also calculated a relative distance metric (*Z_dc_*) that compares the absolute distance *d_c_*(*S, T*) between a disease and a polyphenol with a reference distribution describing the random expectation. The reference distribution corresponds to the expected distances between two randomly selected groups of proteins matching the size and degrees of the original disease proteins and polyphenol targets in the network. It was generated by calculating the proximity between these two randomly selected groups across 1,000 iterations. The mean *μ_d_* (*S,T*) and standard deviation *σ_d_* (*S,T*) of the reference distribution were used to convert the absolute distance *d_c_* into the relative distance *Z_dc_*, defined as:

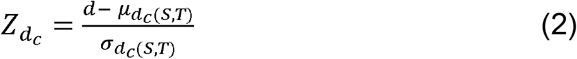

We performed a degree-preserving random selection, but due to the scale-free nature of the human interactome, we avoid repeatedly choosing the same (high degree) nodes by using a binning approach in which nodes within a certain degree interval were grouped together such that there were at least 100 nodes in the bin. The Supplementary Data 2 reports the proximity scores *d_c_* and 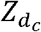 for all pairs of diseases and polyphenols.

### Area Under ROC Curve Analysis

For each polyphenol, we used AUC to evaluate how well the network proximity distinguishes diseases with known therapeutic associations from all of the others of the set of 299 diseases. The set of known associations (therapeutic) retrieved from CTD were used as positive instances, all unknown associations were defined as negative instances, and the area under the ROC curve was computed using the implementation in the Scikit-learn Python package. Furthermore, we calculated 95% confidence intervals using the bootstrap technique with 2,000 resamplings with sample sizes of 150 each. Considering that AUC provides an overall performance, we also searched for a metric to evaluate the top ranking predictions. For this analysis, we calculated the precision of the top 10 predictions, considering only the polyphenol-disease associations with relative distance *Z_dc_* < −0.5^16^.

### Analysis of Network Proximity and Gene Expression Deregulation

We retrieved perturbation signatures from the Connectivity Map database (https://clue.io/) for the MCF7 cell line after treatment with 21 polyphenols. These signatures reflect the perturbation of the gene expression profile caused by the treatment with that particular polyphenol relative to a reference population, which comprises all other treatments in the same experimental plate^26^. For polyphenols having more than one experimental instance (time of exposure, cell line, dose), we selected the one with highest distil_cc_q75 value (75th quantile of pairwise spearman correlations in landmark genes, https://clue.io/connectopedia/perturbagen\_types\_and\_controls). We performed Gene Set Enrichment Analysis^61^ to evaluate the enrichment of disease genes among the top deregulated genes in the perturbation profiles. This analysis offers Enrichment Scores (ES) that have small values when genes are randomly distributed among the ordered list of expression values and high values when they are concentrated at the top or bottom of the list. The ES significance is calculated by creating 1,000 random selection of gene sets with the same size as the original set and calculating an empirical p-value by considering the proportion of random sets resulting in ES smaller than the original case. The p-values were adjusted for multiple testing using the Benjamini-Hochberg method. The network proximity *d_c_* of disease proteins and polyphenol targets for diseases with significant ES were compared according to their therapeutic and unknown-therapeutic associations using the Student’s t-test. The relevant code for calculating the network proximity, AUCs, and enrichment scores can be found on https://github.com/italodovalle/polyphenols.

### Platelet Isolation

Human blood collection was performed as previously described in accordance with the Declaration of Helsinki and ethics regulations with Institutional Review Board approval from Brigham and Women’s Hospital (P001526). Healthy volunteers did not ingest known platelet inhibitors for at least 10 days prior. Citrated whole blood underwent centrifugation with a slow brake (177 x g, 20 minutes), and the PRP fraction was acquired for subsequent experiments. For washed platelets, PRP was incubated with 1 μM prostaglandin E_1_ (Sigma, P5515) and immediately underwent centrifugation with a slow brake (1000 x g, 5 minutes). Platelet-poor plasma was aspirated, and pellets resuspended in platelet resuspension buffer (PRB; 10 mM Hepes, 140 mM NaCl, 3 mM KCl, 0.5 mM MgCl_2_, 5 mM NaHCO_3_, 10 mM glucose, pH 7.4).

### Platelet Aggregometry

Platelet aggregation was measured by turbidimetric aggregometry as previously described^62^. Briefly, PRP was pretreated with RA for 1 hour before adding 250 μL to siliconized glass cuvettes containing magnetic stir bars. Samples were placed in Chrono-Log^®^ Model 700 Aggregometers before the addition of various platelet agonists. Platelet aggregation was monitored for 6 minutes at 37°C with a stir speed of 1000 rpm and the maximum extend of aggregation recorded using AGGRO/LINK^®^8 software. In some cases, dense granule release was simultaneously recorded by supplementing samples with Chrono-Lume^®^ (Chrono-Log^®^, 395) according to the manufacturer’s instructions.

### Platelet Alpha Granule Secretion and Integrin α_IIb_β_3_ Activation

Changes in platelet surface expression of P-selectin (CD62P) or binding of Alexa Fluor^TM^ 488-conjugated fibrinogen were used to assess alpha granule secretion and integrin α_IIb_β_3_ activation, respectively. First, PRP was pre-incubated with RA for 1 hour, followed by stimulation with various platelet agonists under static conditions at 37°C for 20 minutes. Samples were then incubated with APC-conjugated anti-human CD62P antibodies (BioLegend^®^, 304910) and 100 μg/mL Alexa Fluor^TM^ 488-Fibrinogen (Thermo Scientific^TM^, F13191) for 20 minutes before fixation in 2% [v/v] paraformaldehyde (Thermo Scientific^TM^, AAJ19945K2). Fifty thousand platelets were processed per sample using a Cytek^TM^ Aurora spectral flow cytometer. Percent-positive cells were determined by gating on fluorescence intensity compared to unstimulated samples.

### Platelet Cytotoxicity

Cytotoxicity were tested by measuring lactate dehydrogenase (LDH) release by permeabilized platelets into the supernatant^63^. Briefly, washed platelets were treated with various concentrations of RA for 1 hour, before isolating supernatants via centrifugation (15,000 x g, 5 min). A Pierce LDH Activity Kit (Thermo Scientific^TM^, 88953) was then used to assess supernatant levels of LDH.

### Immunoprecipitation and Western blot

Washed platelets were pre-treated with RA for 1 hour, followed by a 15 minute treatment with Eptifibatide (50 μM). Platelets were then stimulated with various agonists for 5 minutes under stirring conditions (1000 rpm, 37°C). Platelets were lysed on ice with RIPA Lysis Buffer System^®^ (Santa Cruz^®^, sc-24948) and supernatants clarified via centrifugation (15,000 x g, 10 min, 4°C). For immunoprecipitation of FYN, lysates were first precleared of IgG by incubating with Protein A agarose beads (Cell Signaling Technologies, 9863S) for 30 minutes at 4°C, before isolation of the supernatant via centrifugation (15,000 x g, 10 min, 4°C). Supernatants were incubated with anti-FYN antibodies (Abcam, 2A10) overnight at 4°C before incubation with Protein A beads for 1 hour. Beads were then washed 5 times with NP-40 lysis buffer (144 mM Tris, 518 mM NaCl, 6 mM EDTA, 12 mM Na_2_VO_3_, 33.3% [v/v] NP-40, Halt^TM^ protease inhibitor cocktail (Thermo, 78429)).

For Western Blot analysis, total cell lysates or immunoprecipitated FYN were reduced with Laemmli Sample Buffer (Bio-Rad, 1610737) and proteins separated by molecular weight in PROTEAN TGX^TM^ precast gels (Bio-Rad, 4561084). Proteins were transferred to PVDF membranes (Bio-Rad, 1620174) and probed with either 4G10 (Milipore, 05-321), a primary antibody clone that recognizes phosphorylated tyrosine residues, or primary antibodies that probe for the site-specific phosphorylation of src family kinases (SFKs, p-Tyr416) within their activation loop. Membranes were incubated with horseradish peroxidase-conjugated secondary antibodies (Cell Signaling Technologies, 7074S) to catalyze an electrochemiluminescent reaction (Thermo Scientific^TM^, PI32109). Membranes were visualized using a Bio-Rad ChemiDoc Imaging System and densitometric analysis of protein lanes conducted using ImageJ (NIH, Version 1.52a).

## Author Contributions

I.F.V and A.L.B designed the study. I.F.V. performed all computational analyses. H.G.R, M.W.M., E.B., and J.L designed and performed experimental validation. J.L. guided I.F.V. on validation case studies. S.M and D.B guided I.F.V for data interpretation and curation of disease associations obtained from literature. I.F.V and A.L.B wrote the paper with input from all authors. All authors read and approved the manuscript.

## Acknowledgements

This study was supported, in part, by NIH grant 1P01HL132825, HG007690, HL108630, and HL119145; American Heart Association grants 151708 and D700382; and ERC grant 810115-DYNASET. We would like to thank Peter Ruppert, Giulia Menichetti, and Istvan Kovacs for support in this study, Feixiong Cheng for assembling the Human Interactome, and Alice Grishchenko for help with data visualization.

## Data Availability

The authors declare that all data supporting the findings of this study are available at https://github.com/italodovalle/polyphenols and within the paper and its supplementary information files.

## Code Availability

Computer code is available at https://github.com/italodovalle/polyphenols

