## Supplementary Information for "Network Medicine Framework Shows Proximity of Polyphenol Targets and Disease Proteins is Predictive of the Therapeutic Effects of Polyphenols"

**Supplementary Notes**

Disease Annotations**:** In this study, we retrieved disease annotations from the Comparative Toxicogenomics Database (CTD), which classify disease-chemical associations as “Therapeutic” (T) or “Marker/Mechanism” (M). M refers to a chemical that correlates with the disease or may act in its aetiology, whereas T indicates that the chemical has a known or a potential therapeutic role in the condition. We identified 2,175 T and 293 M annotations for the polyphenols evaluated in this study and we focused our analysis mainly on the T annotations. We find, however, that when we combined the T and M annotations the predictive performance remains unchanged for most polyphenols (Supplementary Figure 3), helping us add 6 additional polyphenols to the original list of 65 used in the original analysis. This observation suggests that network proximity might capture not only therapeutic effects but any modulatory effect on disease processes, but to validate the precise rule of the M group, further analysis with more extensive data on disease annotations would be necessary. We also highlight that network proximity should be used as prioritization step and other factors should be considered when evaluating real systems, since differences in dosage and time of treatment can be determinant of the beneficial or detrimental effects of polyphenols on diseases^1,2^.

Unveiling the Mechanisms Responsible for the Therapeutic Effects of Specific Polyphenols: To demonstrate how the network-based framework can facilitate the mechanistic interpretation of the therapeutic effects of selected polyphenols, we next focus on vascular diseases (V). Of 65 polyphenols evaluated in this study, we found 27 to have associations to V, as their targets were within the V network neighborhood (Supplementary Table 3). We, therefore, inspected the targets of 15 of the 27 polyphenols with 10 or less targets, as experimentally validating the mechanism of action among the interactions of more than 10 targets would provide complexities beyond the scope of this study. As we discuss next, the network analysis identified direct links between biological processes related to vascular health and the targets of three polyphenols, gallic acid, rosmarinic acid, and 1,4-naphthoquinone (Supplementary Figure 12).

Gallic acid has a single target, SERPINE1, which is also a V-associated protein, resulting in $d_{c}=0$ and $Z_{dc}= -3.02$. An inspection of the LCC formed by V proteins also reveals that SERPINE1 directly interacts with the VD proteins PLG, LRP1, and F2 (Supplementary Figure 12), proteins directly or indirectly related to blood clot formation and dissolution. Indeed, recent studies using *in vivo* models report that gallic acid has protective effects on vascular health^3–7^ and that it attenuates platelet aggregation through the protein kinases MAPK and AKT^8^.

1,4-naphthoquinone targets four proteins, MAP2K1, MAOA, CDC25B, and IDO1, which are proximal to V-associated proteins ($d_{c}=1.25, Z_{d_{c}}$ =$-1.51$) (Supplementary Figure 12). The polyphenol might influence biological processes related to vascular diseases through the action of its target MAP2K1, a gene involved in signaling pathways related to vascular smooth cell contraction and VEGF signaling, and which itself also interacts with 5 V associated proteins (Supplementary Figure 12). Indeed, the effect of 1,4-naphthoquinone and its derivatives on VEGF signaling and angiogenesis has been previously demonstrated^9,10^.

Rosmarinic acid (RA) bind to three human proteins, FYN, MCL1, and AKR1B1, proximal to VD genes ($d_{c}=1.00, Z_{d_{c}}$ =$-1.38$). FYN interacts with three proteins in the V module CD36, APP, and PRKCH) suggesting a role of RA in platelet function, specialized blood cells involved in clot formation and also involved in abnormal clotting or thrombosis. FYN also directly interacts with NFE2L2 (also known as NRF2), a transcription factor that regulates several genes with antioxidant properties. Recent reports show that mice lacking FYN have reduced platelet activity^11–13^ and that RA’s protective effects on vascular calcification and on aortic endothelial function after diabetes-induced damage is mediated by antioxidant mechanisms^14,15^. These observations suggest that RA activity might be mediated by FYN, ultimately regulating the processes of platelet activity and expression of antioxidant genes.

In summary, our analysis suggests that gallic acid activity involves thrombus dissolution processes, rosmarinic acid acts on platelet activation and antioxidant pathways through FYN and its neighbors, and 1,4-naphthoquinone acts on signaling pathways of vascular cells through MAP2K1 activity.

Supplementary Figures

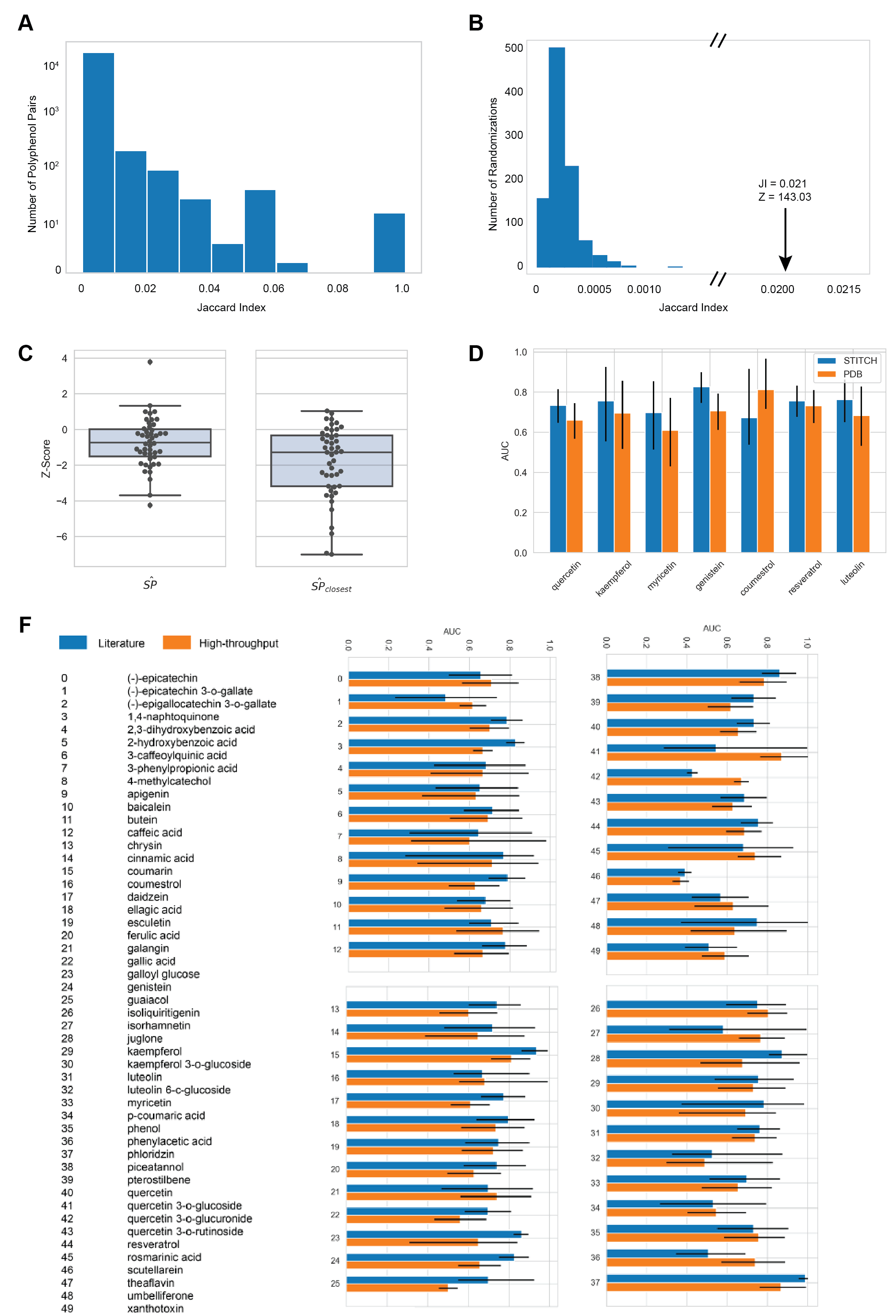

**Supplementary Figure 1 – Target Similarity Among Polyphenols, Network Proximity Among Polyphenol Targets, and Robustness Checks .** A) Distribution of the similarity (Jaccard Index) of the protein targets among polyphenol pairs. B) Expected values of Jaccard Index (JI) average values if the targets of each polyphenol were randomly assigned from the pool of all network proteins with degrees matching the original set. C) Distribution of the network proximity significance among targets of each polyphenol considering the average shortest path among all targets (SP) and the average shortest path to the nearest target (SP_closest_), showing that the targets tend to be proximal to each other compared with random expectation, and that this proximity is even greater when considering the average of distances to the nearest protein. D) Comparison of predictive performance considering PDB as the source of polyphenol protein interactions data**.** PDB provides binding evidence at the 3D resolution level and we could retrieve proteins for 7 polyphenols in PBD. E) Comparison of predictive performance considering the literature-derived interactome assembled in this study and an interactome derived from an unbiased high-throughput screening^16^. We considered the largest connected component of the high-throughput derived interactome, which consisted of 8,955 proteins and 63,619 protein-protein interactions. 49/65 polyphenols could be mapped in both interactomes, while 16/49 could be mapped only in the literature-derived interactome.

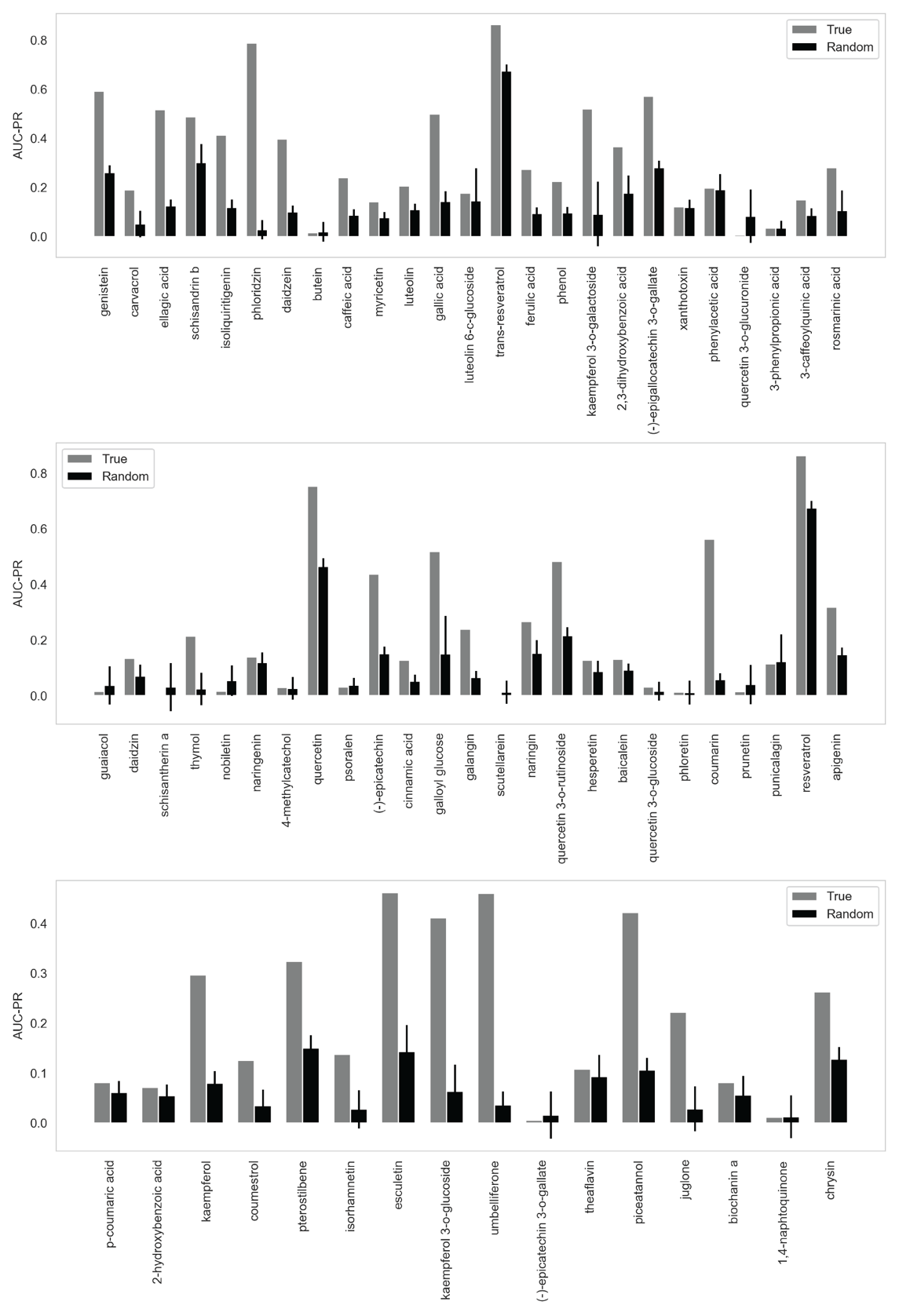

**Supplementary Figure 2 – Predictive Performance Evaluated By The Area Under The Precision Recall Curve (AUC-PR)**. We observed that the AUC-PR is higher than random expectation for most polyphenols evaluated in this study. The random expectation was calculated after randomizing labels across 1000 iterations, being the average and standard deviation represented by the black bars.

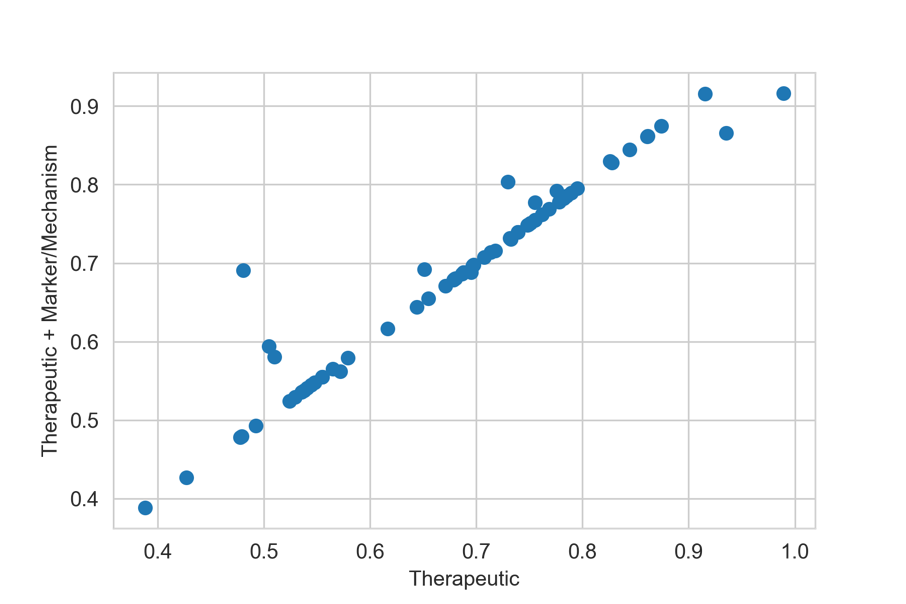

**Supplementary Figure 3 – Comparison of Predictive Performance Using Different Disease Annotation Types.** AUC scores obtained by using different disease annotations from the CTD database. In the vertical axis, the performance was evaluated using disease annotation labels Therapeutic and Marker/Mechanism. In the horizontal axis, the performance was evaluated considering only Therapeutic labels. We highlight that many polyphenol-disease annotations will have both labels, which explains the performance similarity in both cases.

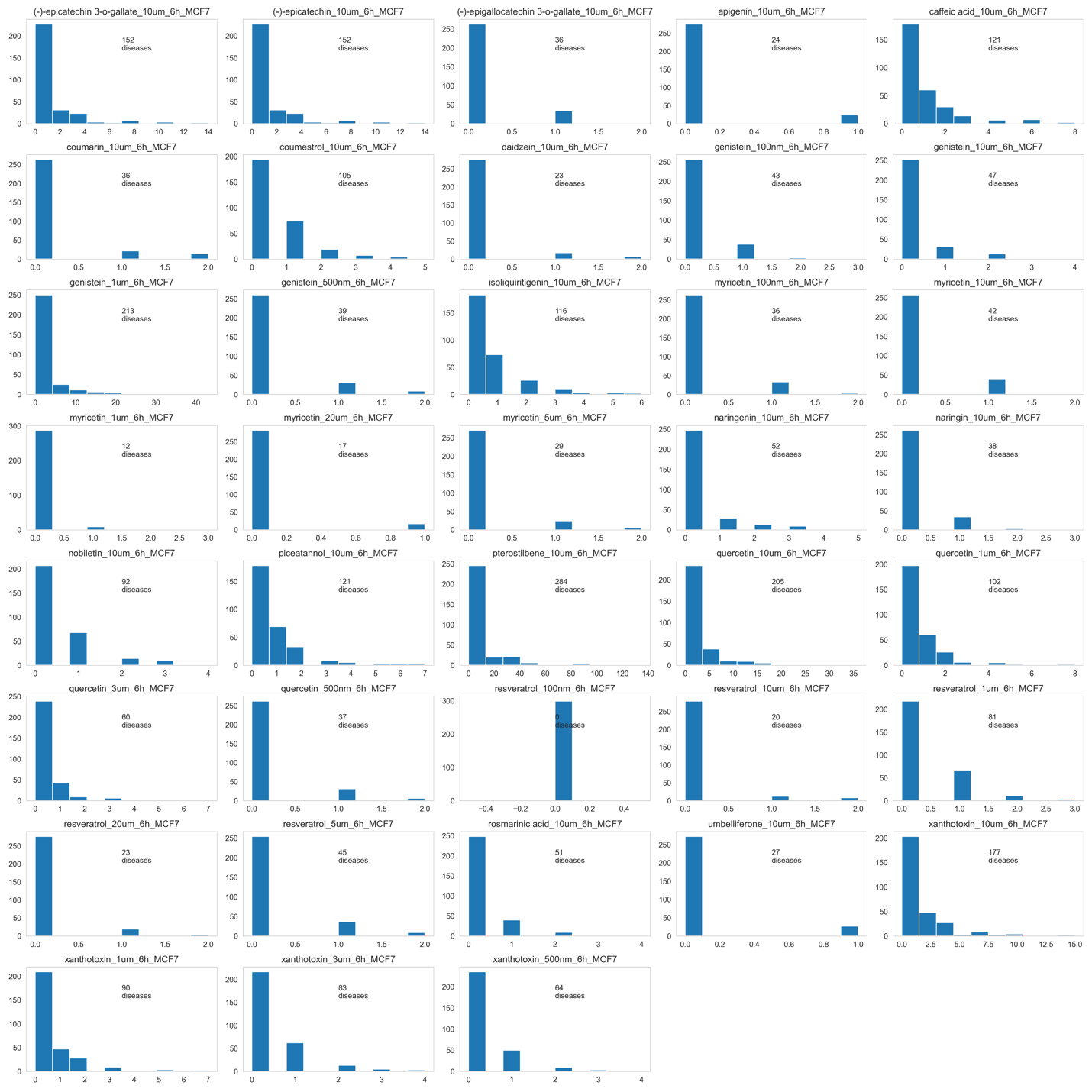

**Supplementary Figure 4 – Distribution of the Number of Perturbed Genes Overlapping With Disease Genes in the Different Experimental Instances**. The horizontal axis of each subplot represents the number of diseases in which disease genes overlap with perturbed genes and the vertical axis represents the frequency of the respective overlap across the 299 diseases. The number of diseases with genes that overlap with at least one perturbed gene is also shown in the plots.

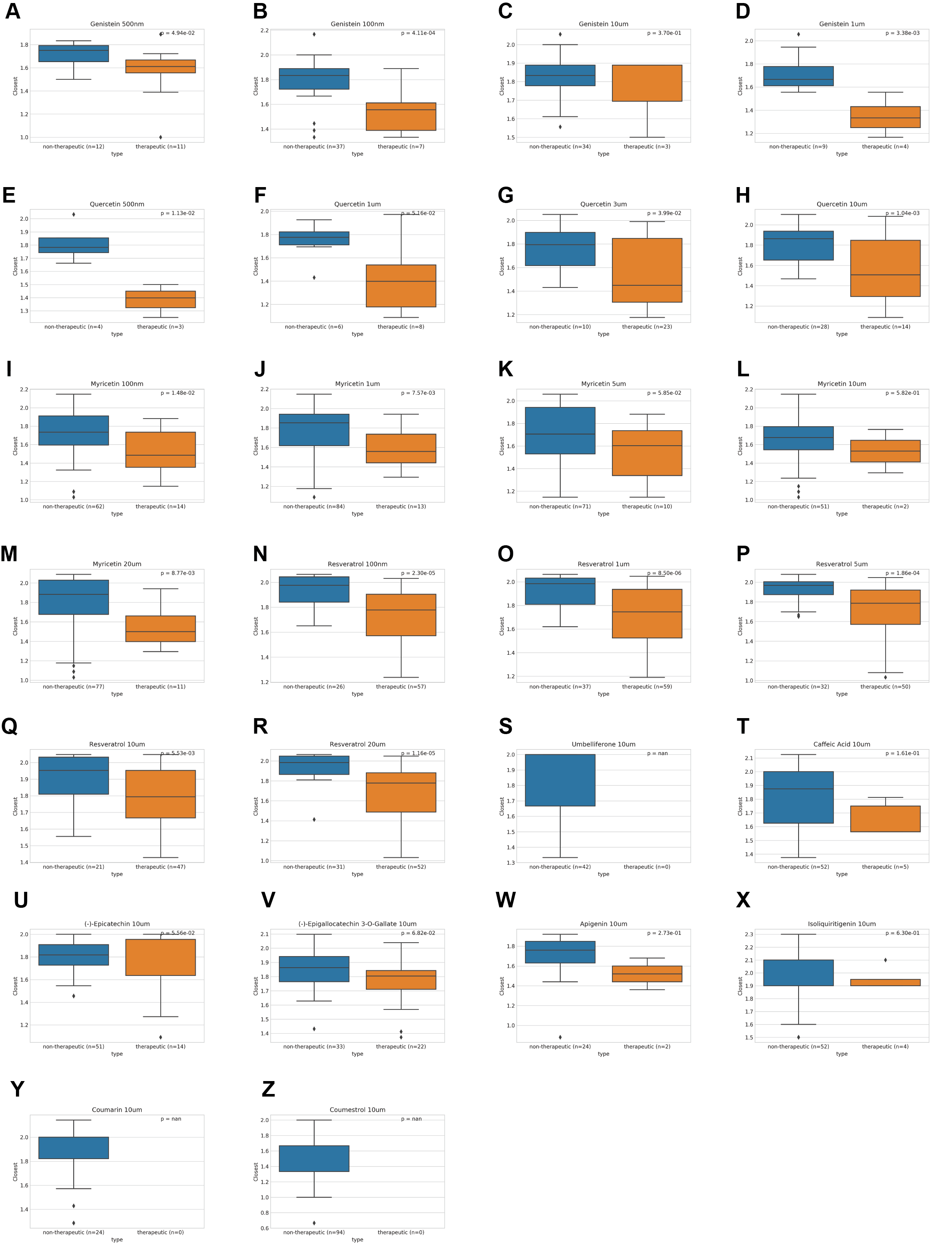

**Supplementary Figure 5 – Enrichment of Perturbated Genes in Expression Profiles Versus Network Proximity.** Among diseases whose genes are enriched with highly perturbed genes, those with therapeutic associations show smaller network distances to the polyphenol targets than those without. A-R) Comparison of polyphenols genistein, quercetin, myricetin, and resveratrol at several concentrations. S-Z) Comparison of polyphenols (-)-epicatechin, (-)-epicatechin 3-O-gallate, caffeic acid, coumarin, coumestrol, daidzein, isoliquiritigenin, and umbelliferone at 10 µM.

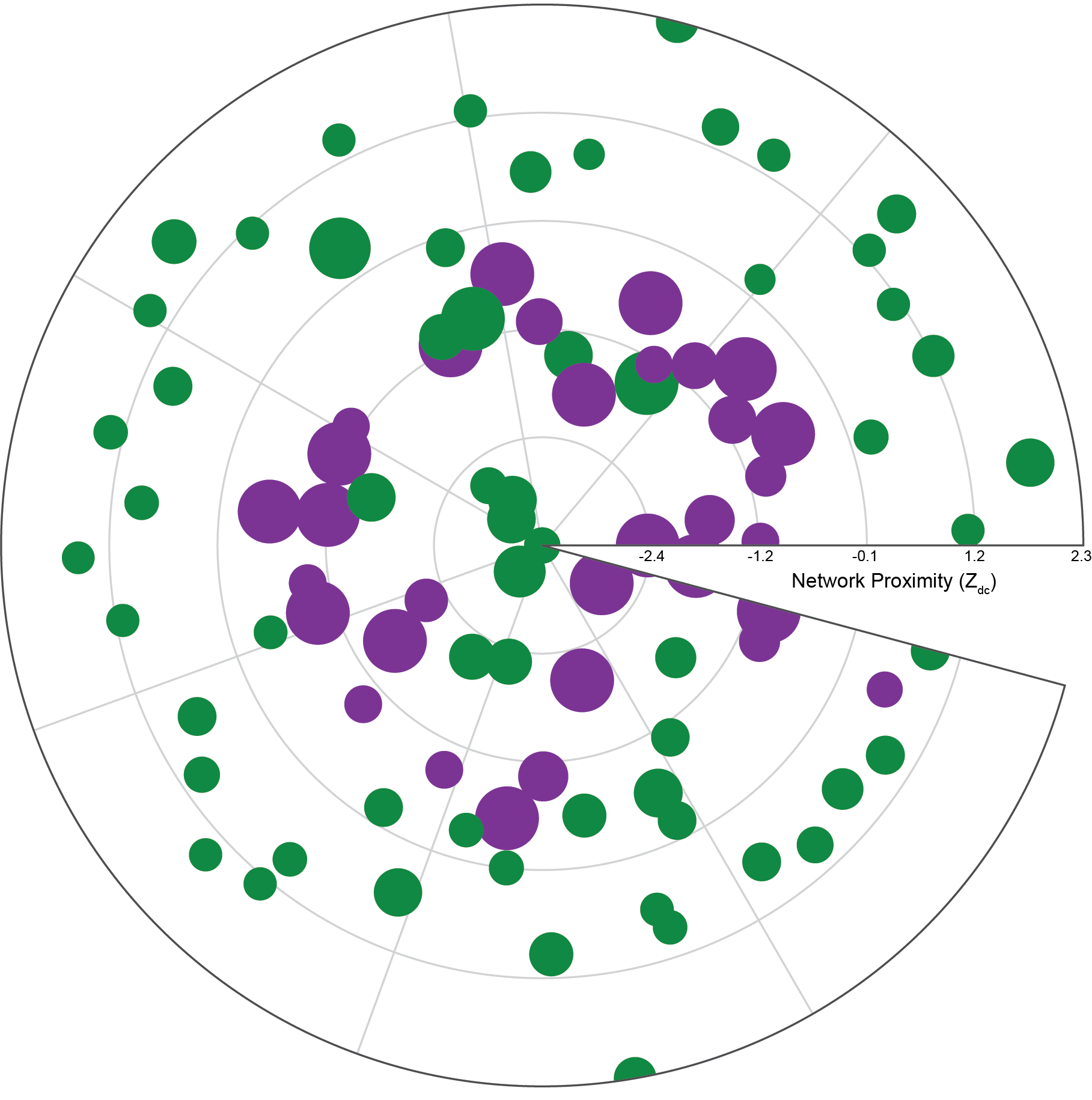

**Supplementary Figure 6 –** **Diseases Proximal to Polyphenol Targets Have Higher Gene Expression Perturbation Profiles.** Proximal and distal diseases in relation to genistein targets. Each node represents a disease and the node size is proportional to the maximum perturbation score after treatment with genistein (1 µM, 6 hours). Distance from the origin represents the network proximity ($Z_{d_{c}}$) to genistein targets. Purple nodes represent diseases in which the therapeutic association was previously known.

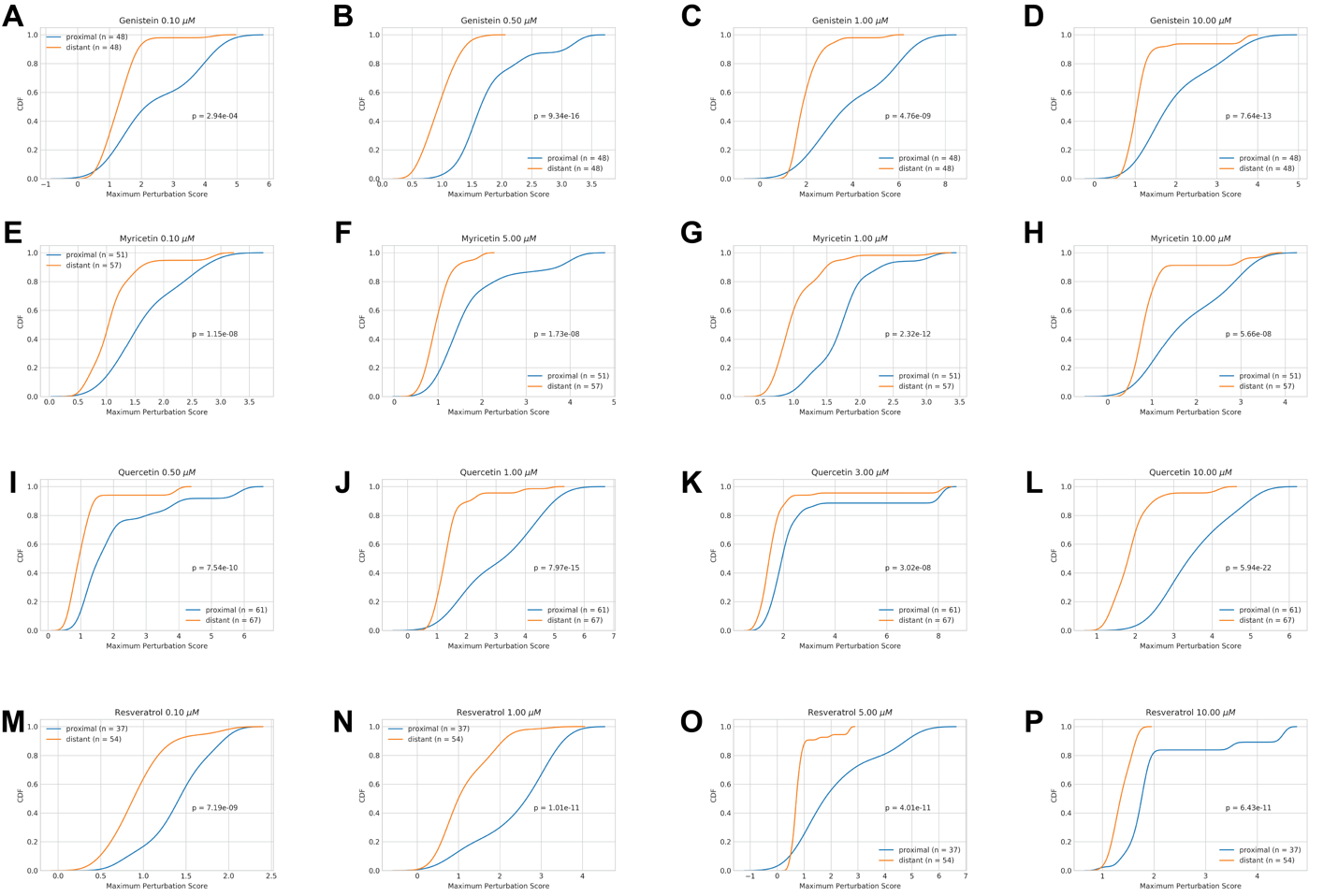

**Supplementary Figure 7 – Diseases Proximal to the Polyphenol Have Higher Perturbation in Expression Profiles of the Cell Line MCF7 Treated with the Respective Polyphenol**. Each disease is represented by the perturbation score of its most perturbed gene in the expression profiles. The comparison of the distribution of proximal and distant diseases was evaluated using the Kolmogorov Smirnov test. Comparisons of polyphenols genistein, myricetin, quercetin, and resveratrol at different concentrations.

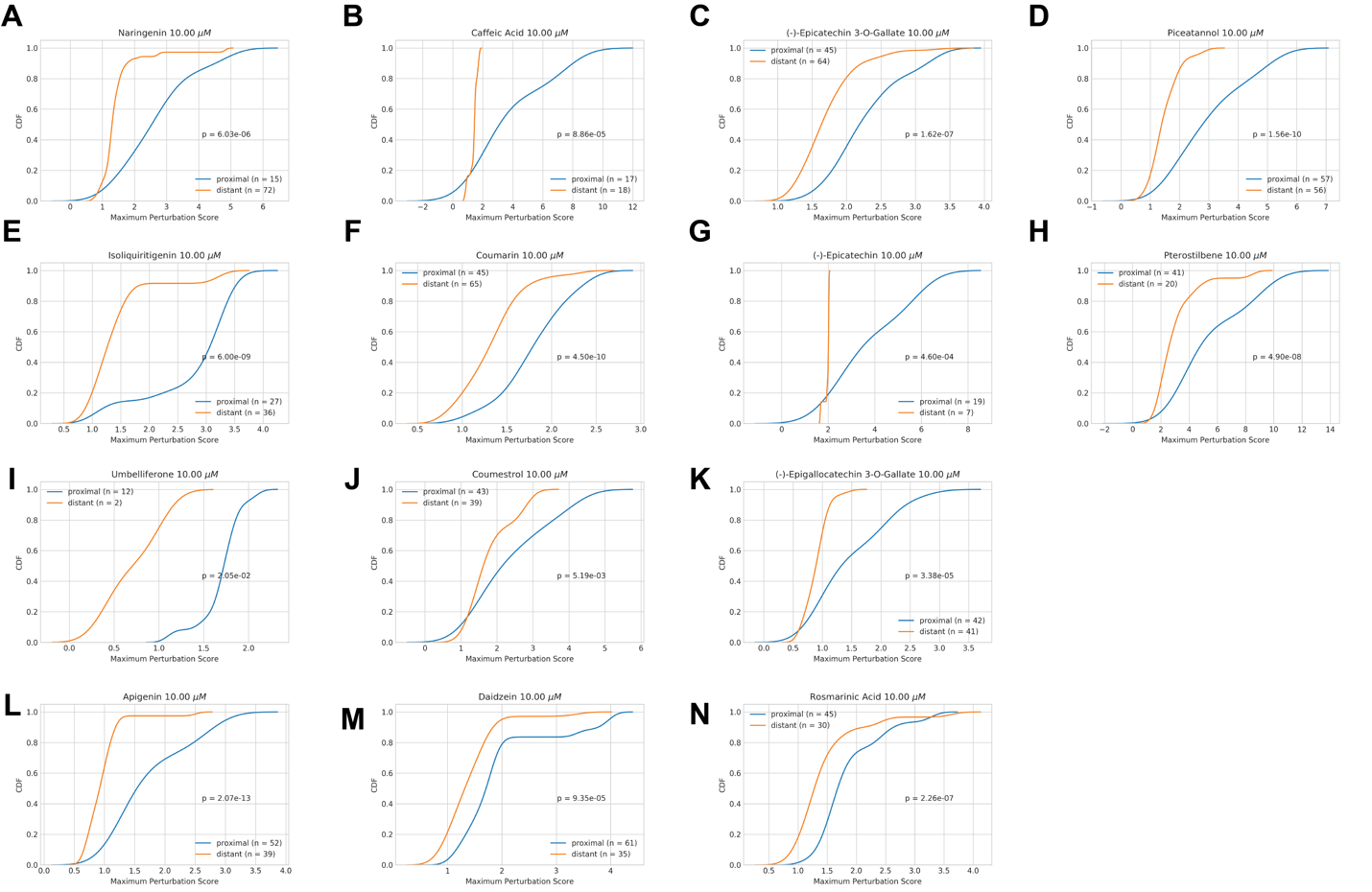

**Supplementary Figure 8 – Diseases Proximal to the Polyphenol Have Higher Perturbation in Expression Profiles of the Cell Line MCF7 Treated with the Respective Polyphenol**. Each disease is represented by the perturbation score of its most perturbed gene in the expression profiles. The comparison of the distribution of proximal and distant diseases was evaluated using the Kolmogorov Smirnov test. Comparisons of polyphenols narigenin, caffeic acid, (-)-epicatechin 3-O-gallate, (-)-epigallocatechin 3-O-gallate, (-)-epicatechin, pterostilbene, piceatannol, apigenin, caffeic acid, coumarin, coumestrol, daidzein, isoliquiritigenin, umbelliferone, and rosmarinic acid at 10 µM.

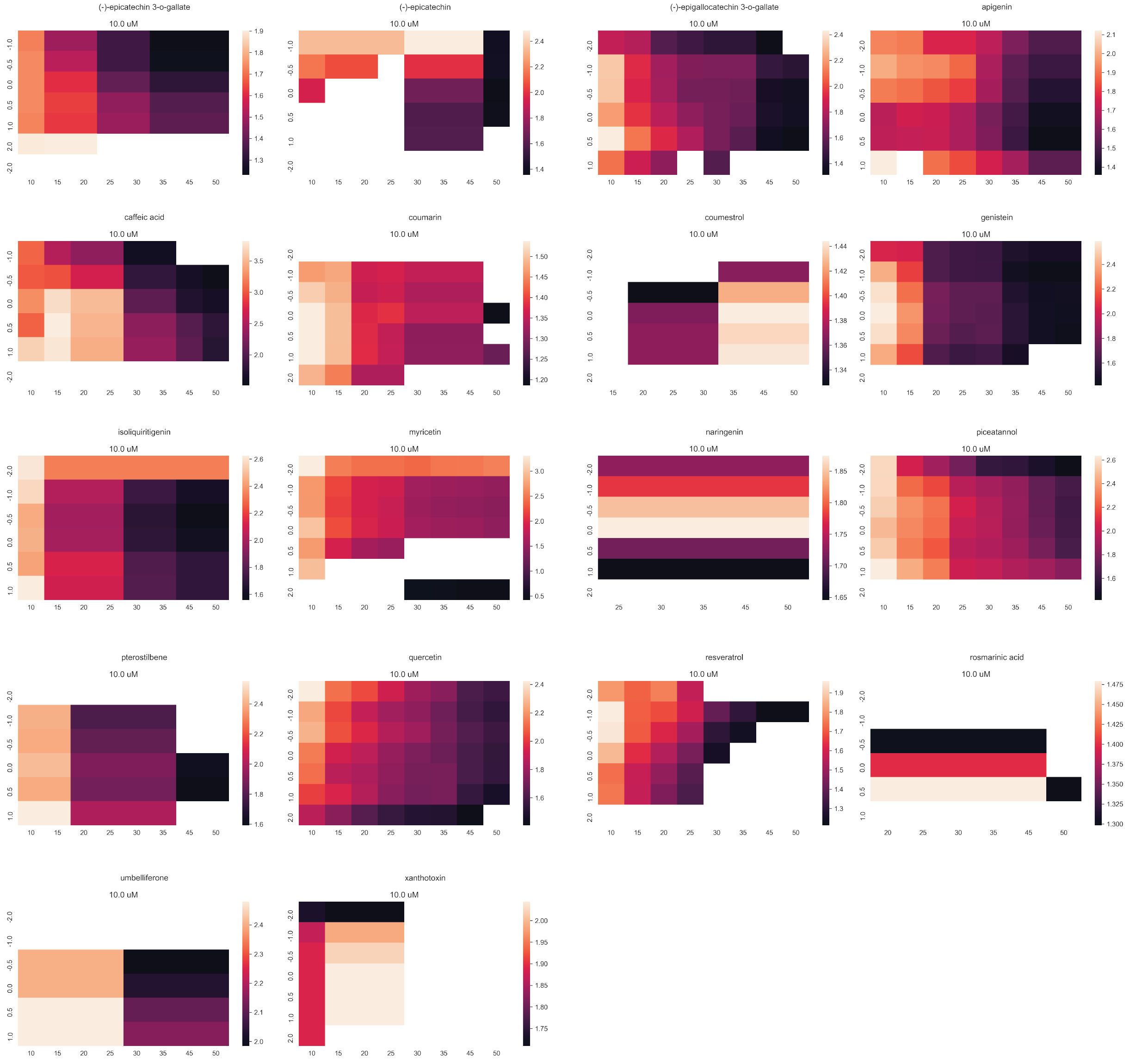

**Supplementary Figure 9 – Test of Different Thresholds When Comparing Gene Expression Perturbations of Proximal and Distant Polyphenols**. The $X$ and $Y$ axes represent different thresholds to determine proximal ($d_{c}< X$ and $Z_{d_{c}}<Y$) and distant ($d_{c}>100- X$ and $Z_{d_{c}}>Y$) diseases. The cells are colored according to the differences in maximum perturbation between proximal and distant diseases (average perturbation proximal / average perturbation distant). White cells represent non-significant differences (T-test > 0.005).

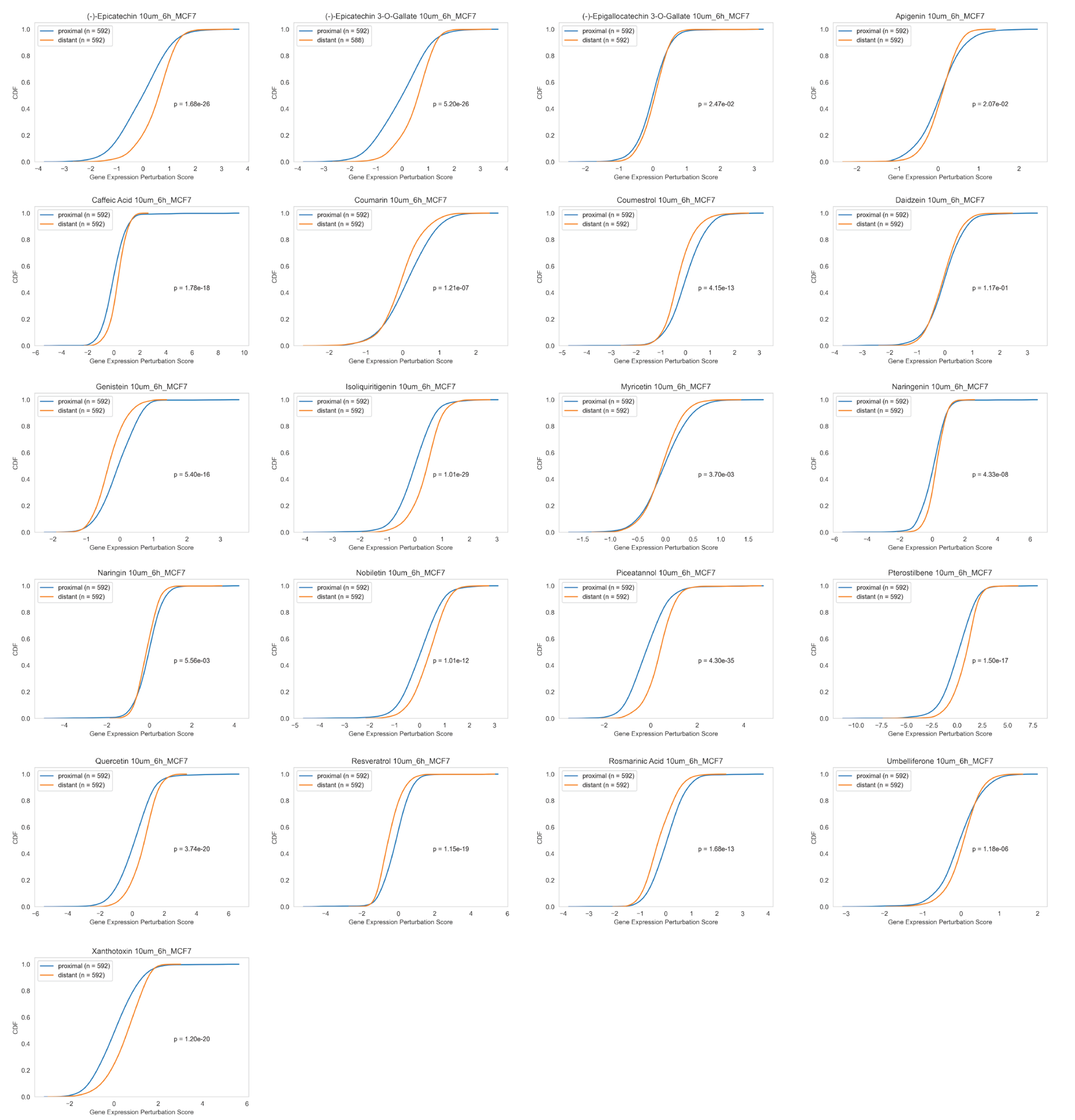

**Supplementary Figure 10 – Gene Expression Perturbations in the Network Vicinity of Polyphenol Targets**. Cumulative Distribution Function (CDF) of the gene expression perturbation scores for distant and proximal proteins in relation to the polyphenol targets. To define proximal and distant proteins, we ranked all proteins in the interactome using a random walk that uses as seed the set of proteins formed by the polyphenol targets. We select the top and bottom 5% genes in terms of the random walk arrival probability to define proximal and distant proteins, respectively. The comparison of the distribution of proximal and distant diseases was evaluated using the Kolmogorov Smirnov test.

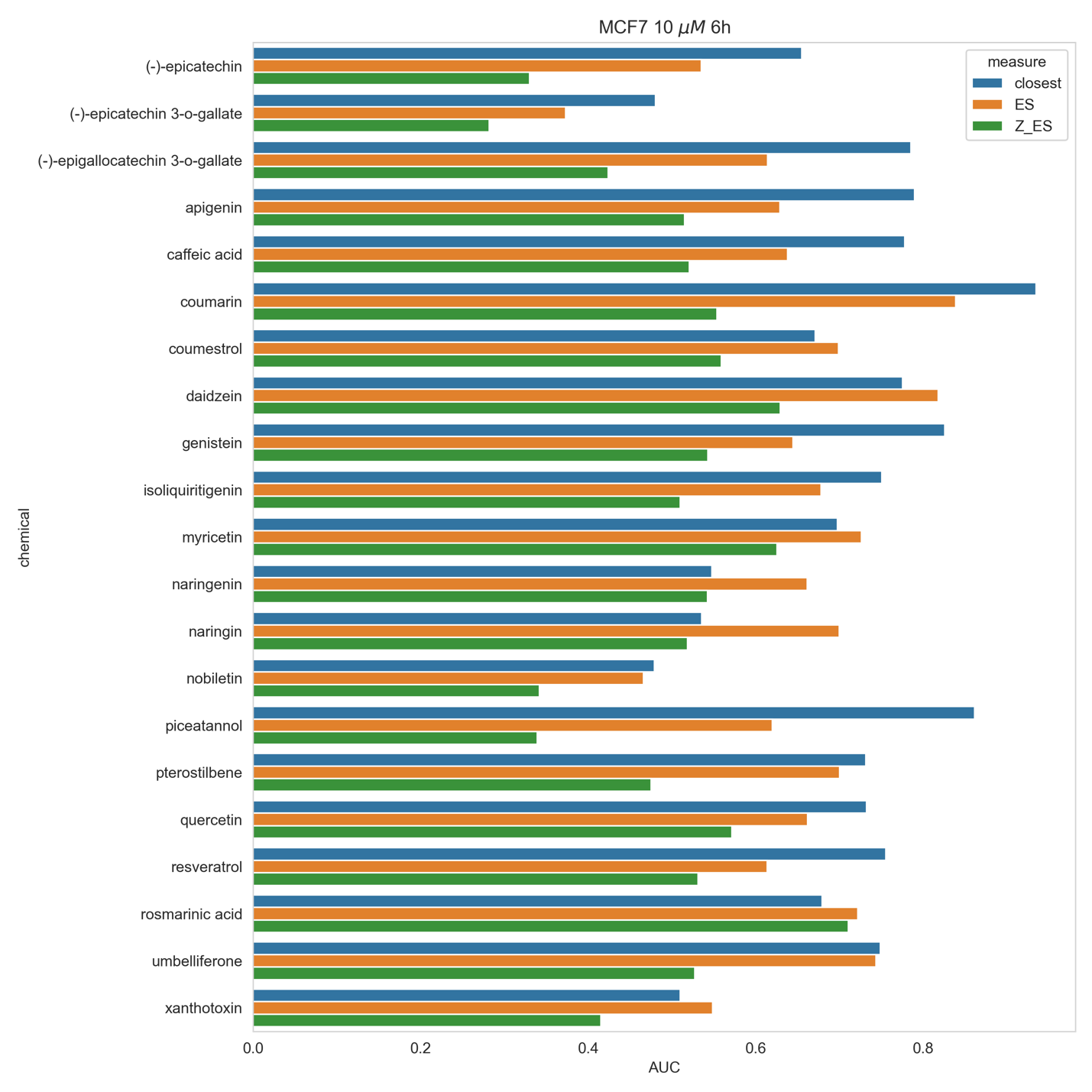

**Supplementary Figure 11 – Comparison of the Predictive Performance When Using Network Proximity or Gene Expression Perturbation Enrichment**. The X axis represent the AUC scores calculated when using network proximity (closest) or the enrichment score in gene expression perturbation profiles (ES, Z_ES). ES: Enrichment Score. Z_ES: z-score calculated comparing the real ES with the average and standard deviation obtained in 1000 permutations with randomly selected proteins.

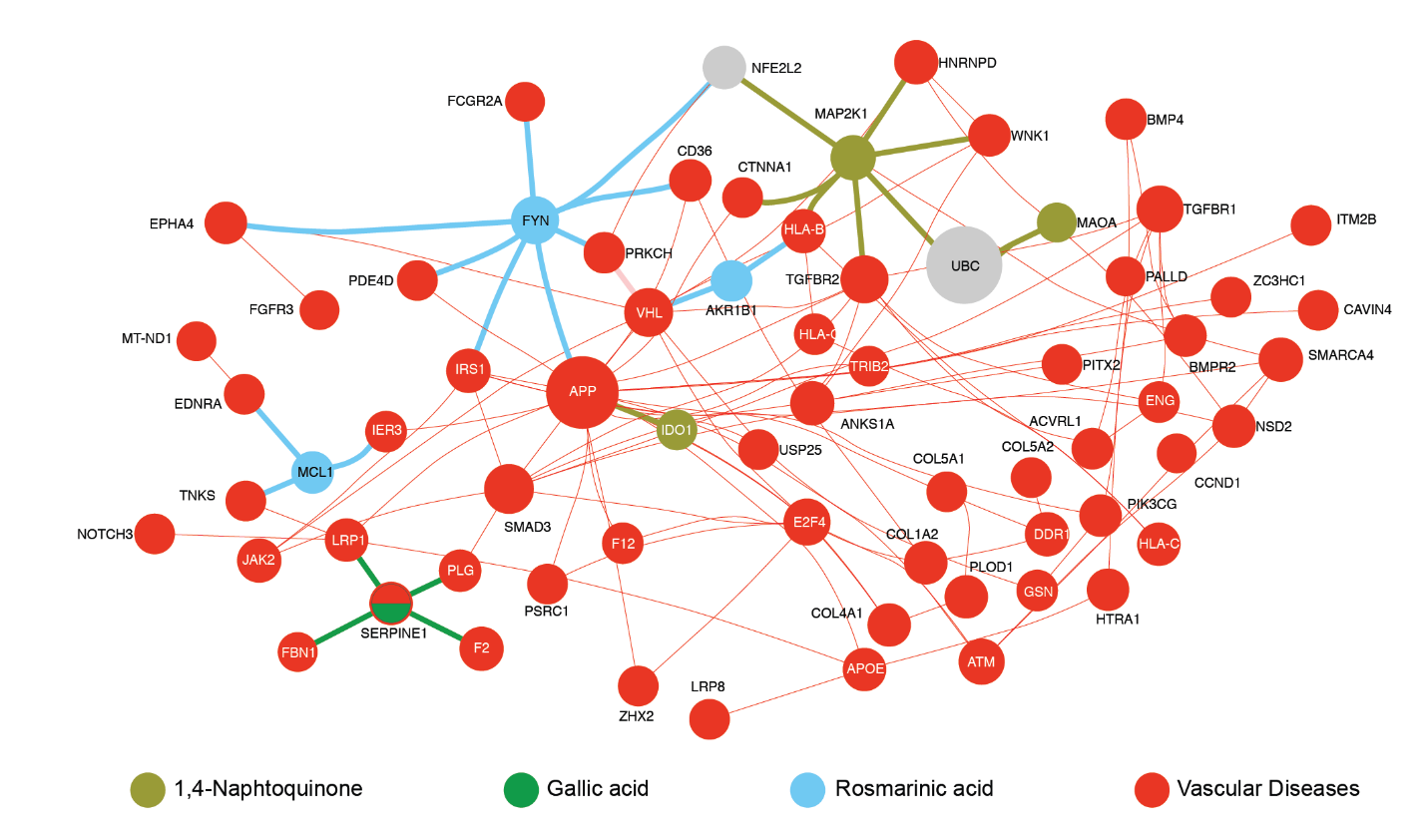

**Supplementary Figure 12 –** **Inferring the Mechanism of Action for Selected Polyphenols.** Interactome neighborhood containing the interactions between proteins associated with vascular diseases and the targets of 1,4-naphthoquinone, gallic acid, and rosmarinic acid**.**

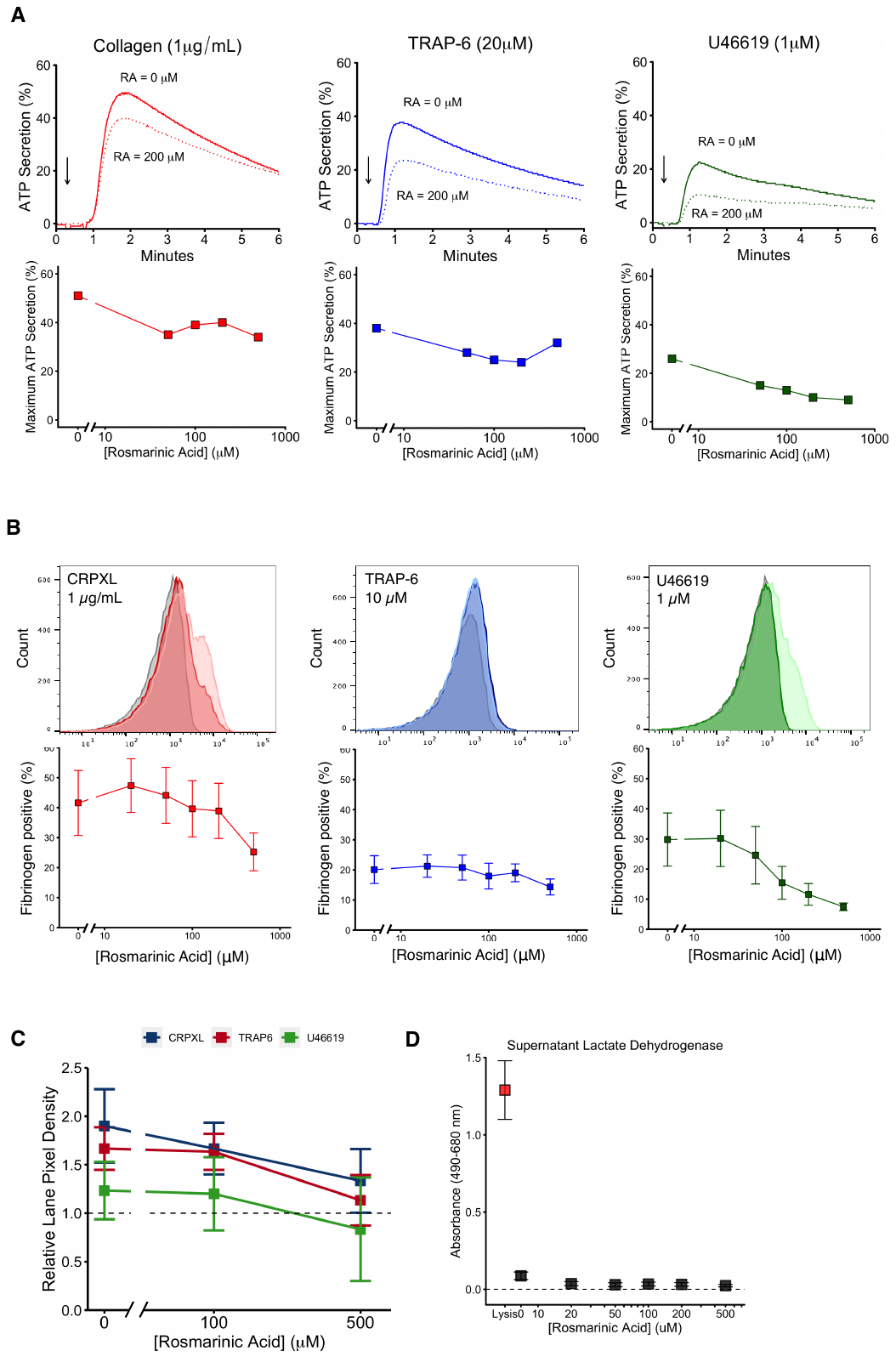

**Supplementary Figure 13	– Rosmarinic Acid Modulates Platelet Dense Granule Release, Integrin Activation and Tyrosine Phosphorylation.** Platelet-rich plasma (PRP) was pre-treated with RA for 1 hour before stimulation with either collagen (1 μg/mL), collagen-related peptide (CRP-XL, 1 μg/mL), thrombin receptor activator peptide-6 (TRAP-6, 10,20 μM), or U46619 (1 μM). Platelets were assessed for either (B) dense granule secretion or (C) integrin α_IIb_β_3_ activation. Arrows indicate the time of agonist addition. Grey histograms represent unstimulated samples, lightly shaded histograms represent samples with no RA pretreatment and filled histograms represent stimulation with prior RA treatment (100 μM). (C) Washed platelets were pre-treated with rosmarinic acid (RA) for 1 hour and supernatants tested for lactate dehydrogenase (LDH). Red box indicates platelets lysed with Triton X-100, dashed line indicates basal LDH release from untreated platelets. (D) Platelet lysates were probed with the antibody 4G10 to measure total tyrosine phosphorylation. N = 1-6 separate blood donations, mean +/- SEM.

**Supplementary Tables**

**Supplementary Table 1 - Summary of Polyphenols Evaluated in this Study.** Name, class, subclass and PubChem IDs for polyphenols. The table also shows the number of polyphenol protein targets mapped in the human interactome, the size of the largest connected component (LCC) formed by them and z-score for the LCC size. The columns min (µM) and max (µM) report the minimum and maximum polyphenol concentrations detected in blood according to Human Metabolome Database (HMDB).

**Supplementary Table 2 - Predicted Gastrointestinal (GI) Absorption and Bioavailability**. Predictions obtained from the SwissADME webserver. The column ‘bioavailability score’ reports the probability of a compound to have at least 10% oral bioavailability in rat or of having measurable Caco-2 permeability.

**Supplementary Table 3 – Polyphenols Proximal to Vascular Diseases.**

**Supplementary Data**

Supplementary Data 1 – .txt file containing the human interactome assembled in this study.

Supplementary Data 2 – .csv file containing the network proximity calculations between 65 polyphenols and 299 diseases.
